# TALPID3/KIAA0586 regulates multiple aspects of neuromuscular patterning during gastrointestinal development in animal models and human

**DOI:** 10.1101/2021.05.28.446103

**Authors:** Jean Marie Delalande, Nandor Nagy, Conor J. McCann, Dipa Natarajan, Julie E. Cooper, Gabriela Carreno, David Dora, Alison Campbell, Nicole Laurent, Polychronis Kemos, Sophie Thomas, Caroline Alby, Tania Attié-Bitach, Stanislas Lyonnet, Malcolm P. Logan, Allan M. Goldstein, Megan G. Davey, Robert M.W. Hofstra, Nikhil Thapar, Alan J. Burns

## Abstract

TALPID3/KIAA0586 is an evolutionary conserved protein, which plays an essential role in protein trafficking. Its role during gastrointestinal (GI) and enteric nervous system (ENS) development has not been studied previously. Here, we analysed chicken, mouse and human embryonic GI tissues with TALPID3 mutations. The GI tract of TALPID3 chicken embryos was shortened and malformed. Histologically, the gut smooth muscle was mispatterned and enteric neural crest cells were scattered throughout the gut wall. Analysis of the Hedgehog pathway and gut extracellular matrix provided causative reasons for these defects. Interestingly, chicken intra-species grafting experiments and a conditional knockout mouse model showed that ENS formation did not require TALPID3, but was dependent on correct environmental cues. Surprisingly, the lack of TALPID3 in enteric neural crest cells (ENCC) affected smooth muscle and epithelial development in a non cell-autonomous manner. Analysis of human gut fetal tissues with a *KIAA0586* mutation showed strikingly similar findings compared to the animal models demonstrating conservation of TALPID3 and its necessary role in human GI tract development and patterning

## INTRODUCTION

During embryonic development, normal organogenesis depends on tightly orchestrated interactions between cells of different lineages. In the developing gastrointestinal (GI) tract, such interactions occur between ectoderm-derived neural crest cells (NCC) and the embryonic gut, which is derived from lateral mesoderm and endoderm (Zorn and Wells 2009; Noah et al. 2011). Interactions between NCC and the developing gut subsequently determine the functional architecture of the enteric nervous system (ENS) by assuring the correct anatomical localisation of the enteric neuronal plexuses and the establishment of appropriate interconnections with GI smooth muscle and the mucosa (Sasselli et al. 2012; Goldstein et al. 2013; Rao and Gershon 2016; Nagy and Goldstein 2017).

Amongst the numerous proteins that have been shown to be essential for correct vertebrate development is the TALPID3 protein, encoded by the *KIAA0586* gene in human (OMIM 610178). TALPID3 is a ubiquitously expressed protein. Its most recognized function is its requirement for ciliogenesis, as loss of function mutations in the *TALPID3* gene are characterized by a lack of primary cilia in model organisms (Davey et al. 2006; Davey et al. 2007; Bangs et al. 2011; Ben et al. 2011; Davey et al. 2014; Fraser and Davey 2019; Yan et al. 2020). It has been shown that the TALPID3 protein has an evolutionary conserved intracellular localization at the centrosome, which plays a critical role in ciliogenesis and coordination of ciliary protein trafficking, in particular through functional interactions with Rab8 and Mib1 (Yin et al. 2009; Ben et al. 2011; Mahjoub 2013; Sung and Leroux 2013; Villumsen et al. 2013; Kobayashi et al. 2014; Wu et al. 2014; May-Simera et al. 2016; Wang et al. 2016; Li et al. 2017; Naharros et al. 2018). TALPID3 has also been shown to be important for centriole duplication (*via* direct binding to CEP120), as well as centriolar satellite dispersal, centrosome length and orientation, which regulates overall tissue polarity (Wu et al. 2014; Stephen et al. 2015; Tsai et al. 2019). In human, the phenotypic spectrum of *KIAA0586* mutations (the human ortholog of chicken *talpid^3^*) expands from embryonic lethal ciliopathies to paediatric ciliopathy symptoms including Joubert Syndrome (JBTS) (Akawi et al. 2015; Alby et al. 2015; Bachmann-Gagescu et al. 2015; Malicdan et al. 2015; Roosing et al. 2015; Stephen et al. 2015; Alby et al. 2016). A conditional deletion of *talpid^3^* in the central nervous system of a mouse model recapitulates the cerebellar phenotype seen in JBTS (Bashford and Subramanian 2019).

The severe developmental defects caused by the lack of TALPID3 can be linked to the disruption of key developmental signaling pathways, with the strongest association shown to be with the Hedgehog (Hh) pathway (Davey et al. 2006; Davey et al. 2007; Ben et al. 2011; Davey et al. 2014; Ingham 2016; Matsubara et al. 2016; Fraser and Davey 2019). Three Hh gene homologs have been described in vertebrates: *Sonic Hedgehog* (*shh*), *Indian Hedgehog* (*ihh*), and *Desert Hedgehog* (*dhh*) (Pathi et al. 2001; Ingham 2016). At the sub-cellular level, following the binding of Hedgehog ligands to the Patched receptor (PTCH1), the trans-membrane transducer Smoothened (SMO) is transported to the primary cilium by anterograde trafficking. Subsequently, GLI proteins located within the cilium tip are processed into activator (GLIA) or repressor (GLIR) isoforms, which are then released in the cytoplasm. The processing of GLI proteins through the cilium establishes the ratio of GLIA to GLIR proteins, which in turn act as transcriptional effectors to control downstream SHH target genes (Kim et al. 2009; Pan et al. 2009; Sasai and Briscoe 2012; Briscoe and Therond 2013; Ramsbottom and Pownall 2016). TALPID3 has been shown to interact and colocalise with the PKA regulatory subunit PKARIIβ at the centrosome. This interaction leads to the phosphorylation of GLI2 and GLI3 and directly links TALPID3 to a functional step in the Hh pathway (Li et al. 2017).

Many studies have demonstrated the central role of the Hh pathway in gut development, physiology and cancer [reviewed in (Fukuda and Yasugi 2002; van den Brink 2007; Merchant 2012)]. Normal gut development has both common and separate requirements for SHH and IHH. In mouse models, mutations in *shh* or *ihh* result in reduced smooth muscle mass, gut malrotation and annular pancreas (Ramalho-Santos et al. 2000). In addition, *shh* mutants exhibit specific defects such as intestinal transformation of the stomach, duodenal stenosis, increased enteric neurons, abnormally distributed ganglia and imperforate anus. On the other hand, *ihh* mutants show reduced epithelial stem cell proliferation and differentiation rate, as well as aganglionic colon (Ramalho-Santos et al. 2000). Interestingly, mutant mice lacking hedgehog-binding protein growth arrest–specific gene 1 (Gas1) or its intracellular messenger Gnaz, have a shortened digestive tract, reduced smooth muscle mass, increased number of enteric neurons and miss-patterned ENS (Kang et al. 2007; Biau et al. 2013). This phenotype has been attributed to a combination of reduced Hh signaling and increased Ret tyrosine kinase signaling (Biau et al. 2013). The Ret tyrosine kinase is essential for ENS development (Natarajan et al. 2002).

Here, we examined the GI tracts of *talpid^3^* chicken and human fetal gut tissues bearing a homozygous null mutation in *KIAA0586*. We found remarkably similar phenotypes and comparable defects in gut tissues (Schock et al. 2016). We also investigated the role of *TALPID3* in early formation of the ENS, using chicken chimeras and a *talpid^3^* conditional knock out mouse. We demonstrate that *TALPID3* is not required cell autonomously for ENCC migration and early ENS patterning. Rather, our results demonstrate that *TALPID3* is essential for normal spatial differentiation of smooth muscle and proper expression of ECM components. Our study also reveals that growth and development of both mucosa & smooth muscle are regulated by the ENS *via TALPID3*-mediated signalling.

## RESULTS

### *talpid^3^* chicken embryos have multiple anatomical defects including gastrointestinal defects

*talpid^3^* is a naturally occurring chicken mutant. Although limbs and organ defects have been reported in *talpid^3^* chicken embryos, there has been a lack of detailed analysis of the GI defects in these mutants. As previously described, (Buxton et al. 2004) E10.5 *talpid^3^* chick embryos were smaller than controls, showed generalized oedema and displayed a wide range of congenital abnormalities including short ribs, polydactylous paddle shaped limbs, organ defects (lung hypoplasia, liver fibrosis, cholestasis) and craniofacial abnormalities (hypotelorism, reduction and anterior displacement of the frontonasal process) (Fig. 1A). At E10.5, the GI tract of *talpid^3^* chick embryos was significantly reduced in length compared to controls (Fig. 1B). As previously described, no left right asymmetry or rotation defect was observed in the stomach of *talpid^3^* chick embryos (Stephen et al. 2014). Transverse sections at the level of the neck showed abnormal connection (fistula) between the esophagus and the trachea or a total absence or narrowing (atresia) of either of the structures (Fig. 1G,O; Fig. 4E,I). Additionally, *talpid^3^* embryos had an open hindgut (Fig. 1B–inset-,J,R; Fig. 2H,P; Fig. 4H,L). Lastly, gut epithelium thickness varied between controls and mutants, as shown in Fig. 1M,Q insets.

**Figure 1:**
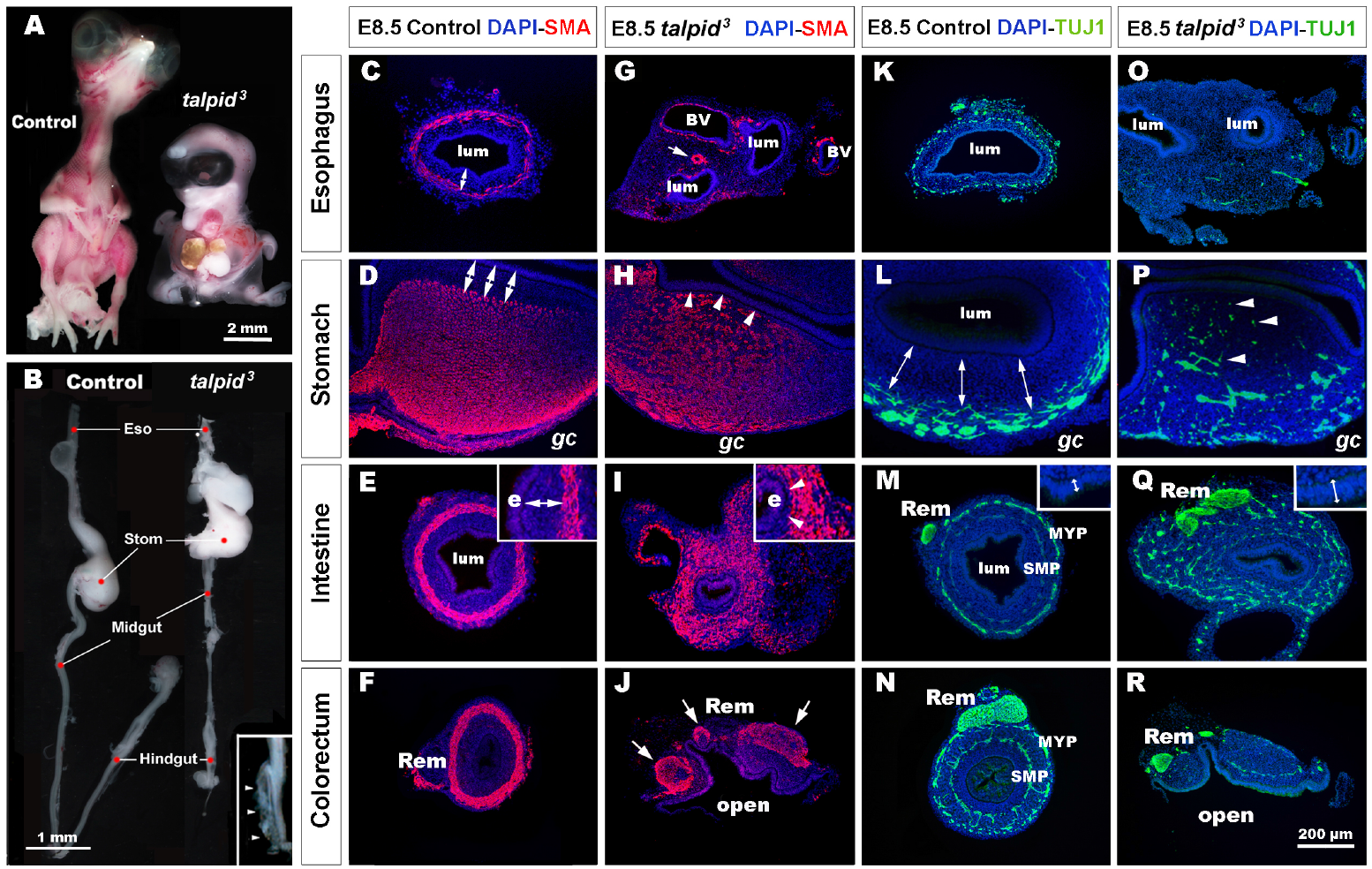
*talpid^3^* embryos have multiple gastrointestinal defects, including disruption of the neuromuscular patterning of the digestive tract. (A) Phenotype of control (left) and *talpid^3^* (right) embryos at E10.5. *talpid^3^* mutant embryos have paddle-shaped limbs and hypotelorism. (B) Anatomical features of the GI tract along the rostro-caudal axis (esophagus, stomach, intestine, caecum, colorectum) of E10.5 control embryos (left). In stage-matched *talpid^3^* embryos (right), all gut regions are apparent, but the esophagus and trachea are malformed, midgut and hindgut are reduced in length, and the terminal hindgut is open. (Inset in B) High magnification of an open hindgut (arrowheads). (C-J) Expression of smooth muscle actin (SMA) at E8.5 in controls (C-F) and (G-J) *talpid^3^* embryos. (C-F) In controls, SMA is expressed in the presumptive circular muscle layer in the outer mesenchyme, distant from the epithelium (C-E, double arrows). (G) In E8.5 *talpid^3^* embryos, the SMA staining of the oesophagus is discontinuous unlike the SMA surrounding blood vessels (arrows). (H,I) SMA staining in the stomach and midgut of mutants is widespread throughout the gut wall, with staining extending adjacent to the epithelium (e) and the serosa (s) (H,I, arrowheads). (J) SMA staining in the colorectum shows disrupted muscle and open hindgut. (K-R) Expression of TuJ-1 labelled enteric neural crest at E8.5 in (K-N) controls and (O-R) *talpid^3^* embryos. (K-N) In E8.5 controls, TuJ-1 staining shows enteric neural crest organised in one (K) or two plexuses (L-N) clearly separated from the mucosa (double arrows). (M-N) The Remak nerve is a single bundle of fibres. (O-R) in *talpid^3^* E8.5 embryos, TuJ-1 staining shows enteric neural crest cells are absent or scattered throughout the mesenchyme (P-R, arrowhead). The Remak nerve is divided in several bundles (Q-R). Epithelium thickness varies between control and mutant intestine (M,Q – inset). m: mucosa; Rem; nerve of Remak; MYP: myenteric plexus; SMP: submuscosal plexus.

**Figure 2:**
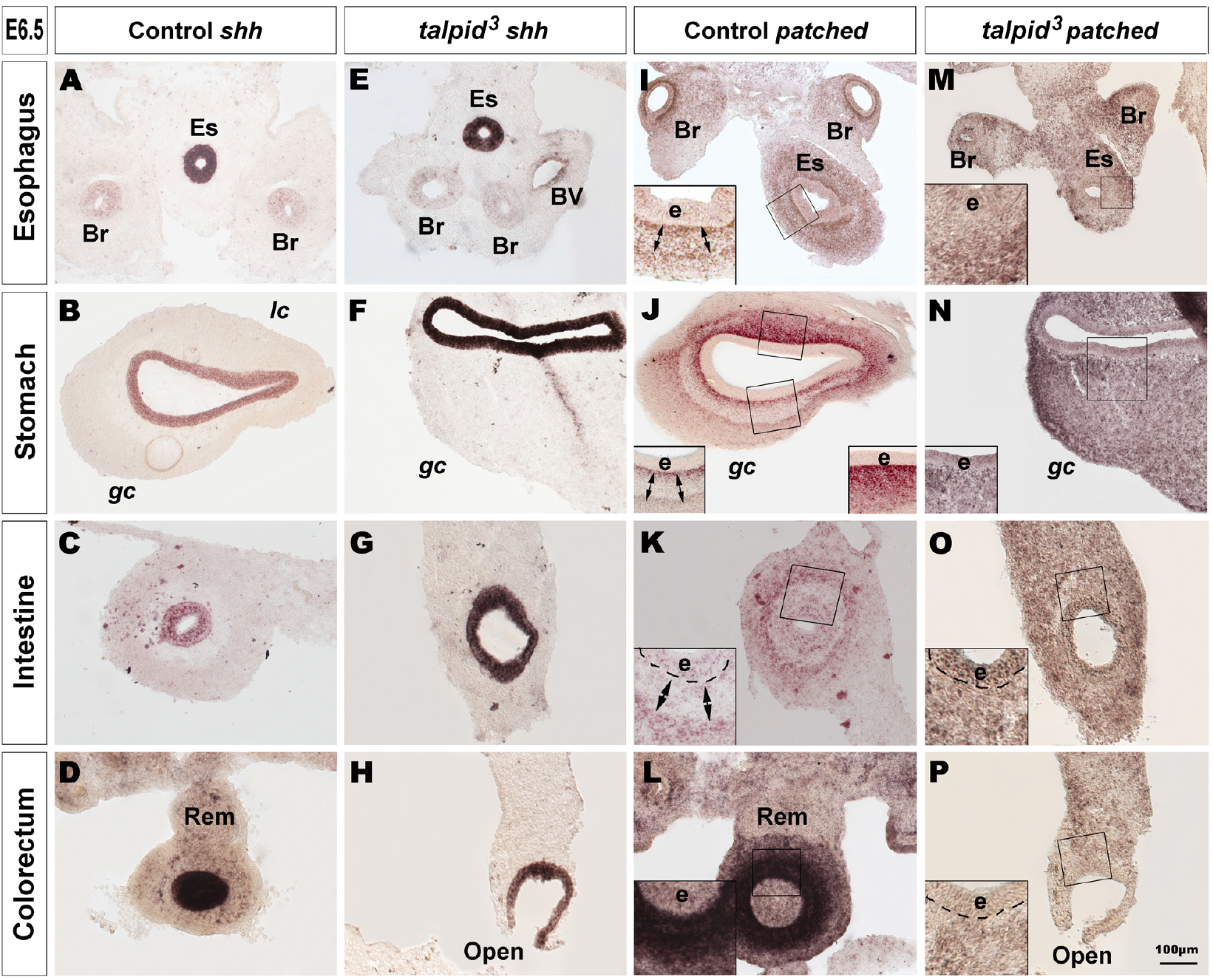
Hh signaling is disrupted in the GI tract of *talpid^3^* mutants. (A-P) *In situ* hybridization for *shh* (A-H) and *patched* (I-P) in E6.5 control (A-D, I-L) and E6.5 *talpid^3^* mutant (E-H, M-P) transverse sections. (A-H) In both controls and *talpid^3^*, *shh* is expressed in the endoderm in all gut regions. (I-L) *Patched in situ* in controls shows a discrete pattern of expression with one (J-L + insets) or two (I-K + Insets) concentric gradients. (M-P + insets) In *talpid^3^* sections the *patched* gradient is replaced by uniform levels of expression. Es: esophagus; Br: bronchus; Rem: nerve of Remak; e: endoderm; *lc*: lesser curvature; *gc*: greater curvature of the stomach.

### Severe smooth muscle and ENS patterning defects in *talpid^3^* chicken GI tract

To gain further insight into the GI defects of *talpid^3^* embryos we performed immunohistochemistry using molecular markers to highlight the neuromuscular organisation of the GI tract in controls and *talpid^3^* mutants. In E8.5 controls, smooth muscle actin (SMA) immunostaining revealed a compact ring of immunopositive cells, corresponding to the presumptive circular muscle layers, encircling the gut epithelium (Fig. 1C-F). This pattern was also evident at other stages of development (Fig. 5I-L) and on sections stained with phalloidin (Fig. 4A-D). Notably, this muscular ring was located in the outer mesenchyme, with a distinct separation between the muscle and the gut epithelium (Fig. 1C-F, double arrows). In *talpid^3^* gut, despite the presence of SMA staining surrounding blood vessels, SMA positive cells were discontinuous or absent around the esophagus (Fig. 1G). This impaired smooth muscle differentiation in the esophagus was also evident by the lack of actin accumulation in *talpid^3^* tissues stained with phalloidin (Fig. 4E,I). In the stomach and the intestine, the compact muscular ring seen in controls was replaced by diffuse staining that extended across the entire gut mesenchyme, with SMA positive cells abutting the gut epithelium (Fig. 1H,I, arrowheads; Fig. 3J; Fig. 4F,J). In the colorectum, the circular muscular pattern was disrupted by the open hindgut phenotype (Fig. 1J, Fig. 4H,L). In E8.5 controls, TuJ-1 immunostaining revealed ENCC-derived neurons arranged in characteristic plexuses (Fig. 1K-N). In the esophagus and the stomach, the presumptive ENS was organised in a single plexus located in the outermost mesenchyme (Fig. 1K,L, double arrows). In the intestine and colorectum, ENCC were organised in two plexuses: the myenteric plexus (MYP) and the submuscosal plexus (SMP) (Fig. 1M,N). The nerve of Remak (Rem), an avian specific nerve derived from sacral ENCC, was also positively labelled (Fig. 1N). In E8.5 TALPID3 embryos, ENCC were present throughout the intestine (Fig. 1Q) and colorectum, although well-defined plexuses were not apparent in the distal gut due to the open hindgut phenotype (Fig. 1R). Although ENCC were observed alongside the esophagus at E6.5 (Fig. 3G; Fig. 4E,I), they were not found around the esophagus at E8.5 and did not form a plexus (Fig. 1O). At this stage, ENCC were present in the stomach and intestine regions, but they failed to organise in plexuses and were scattered throughout the gut wall (Fig. 1P,Q). In the colorectum, the nerve of Remak was also smaller in diameter and/or comprised several bundles (Fig. 1Q,R). Overall, these results showed regions such as the esophagus being devoid of both ENS and smooth muscle in *talpid^3^* mutants, whereas regions such as the ventral stomach and the intestine showed scattered ENS and extended regions of smooth muscle differentiation across the mesenchyme.

**Figure 3:**
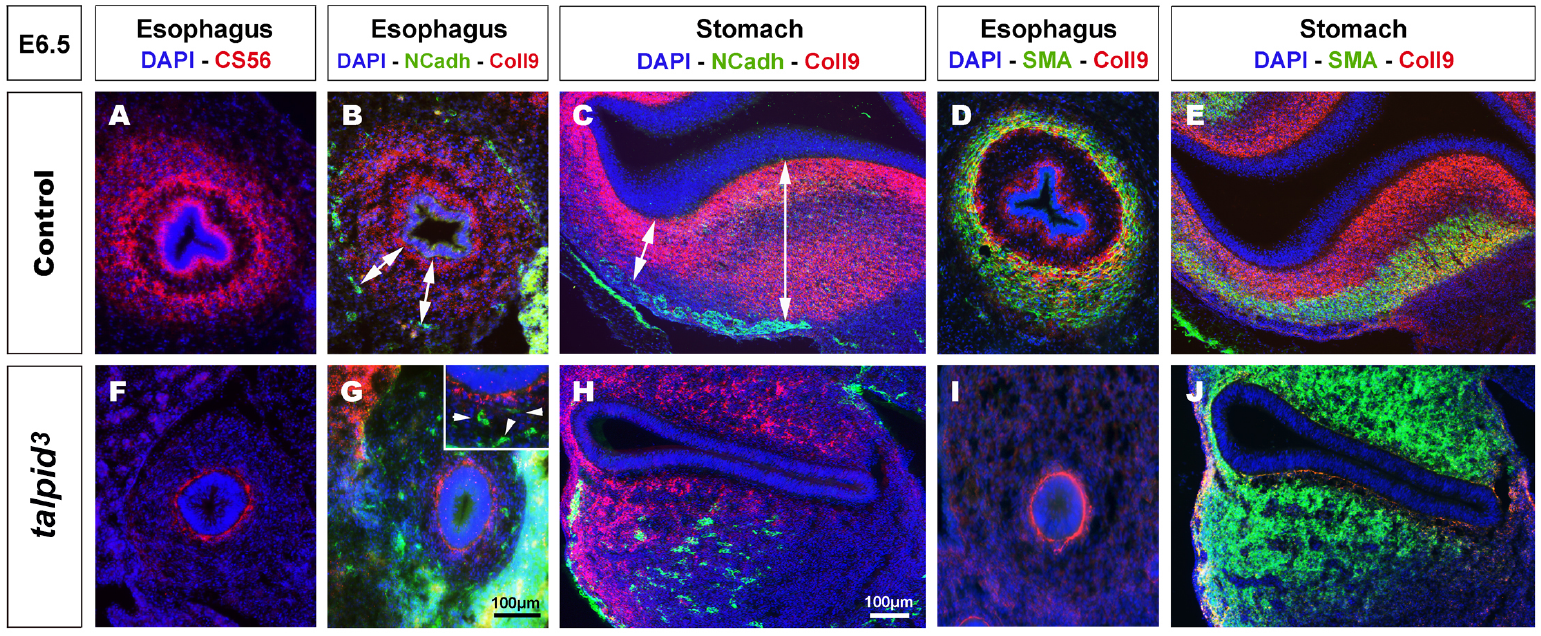
Expression of chondroitin sulfate proteoglycans (CSPG) is altered in *talpid^3^* gut mesenchyme, causing the disappearance of ENCC repellent cues. (A-J) Immunofluorescent staining of esophagus and stomach with CSPG pan marker CS-56 and Coll9, in control (A-E) and *talpid^3^* chicken mutant (F-J) at E6.5, (B,C,G,H) co-stained with N-Cadherin and (D,E,I,J) smooth muscle actin (SMA). (A,B,D) In the esophagus, the pattern of two concentric circles of CSPG expression is lost in (F,G,I) *talpid^3^* mutant (G-inset, arrowheads) coinciding with scattered ENCC. (C) In control stomach, Coll9 expression is widespread and excludes the ENS myenteric plexus whereas in (H, double arrows) *talpid^3^* mutant expression is altered and loss of expression coincides with scattered ENCC. (D,E) In control, SMA expression in partially overlapping with Coll9 (yellow merge). (I,J) in *talpid^3^* mutants, Coll9 expression is altered and SMA is either lost (I) or extends to ectopic sub-epithelial domains (J).

**Figure 4:**
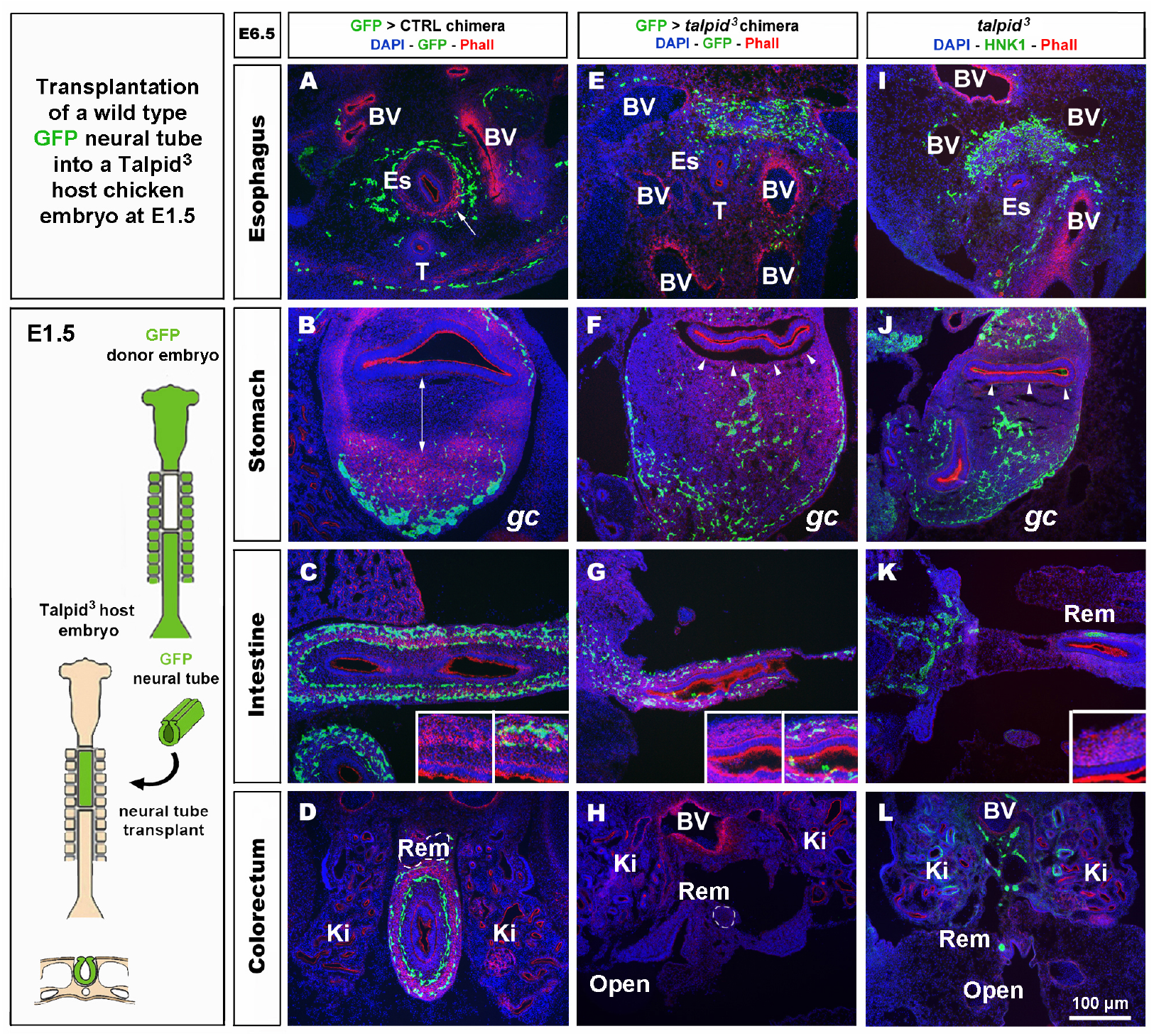
GFP>*talpid^3^* chimeric embryos show ENS and smooth muscle patterning defects similar to stage matched *talpid^3^* embryos. (A-L) E6.5 chimeric embryos after wild type GFP neural tube transplantation in (A-D) GFP>control and (E-H) GFP>*talpid^3^* hosts, compared to (I-L) E6.5 *talpid^3^* embryos. (A-L) Immunofluorescent staining for GFP and Phalloidin in (A-H) chimeric embryos and (I-L) HNK-1 and phalloidin in *talpid^3^* embryos. (E-G + inset) GFP> *talpid^3^* chimeric embryos have scattered ENCC distribution similar to TALPID3 embryos. (A-D) Phalloidin staining shows smooth muscle patterning in control chimera (arrow, double arrow, insets). (E-H) Phalloidin staining shows a lack of smooth muscle patterning in the GFP>*talpid^3^* chimera, similar to the staining found in *talpid^3^* embryos (I-L; arrowheads, inset). BV: blood vessels; Ki: kidneys; Rem: nerve of Remak; Es: esophagus; T: trachea; *gc*: greater curvature of the stomach.

**Figure 5:**
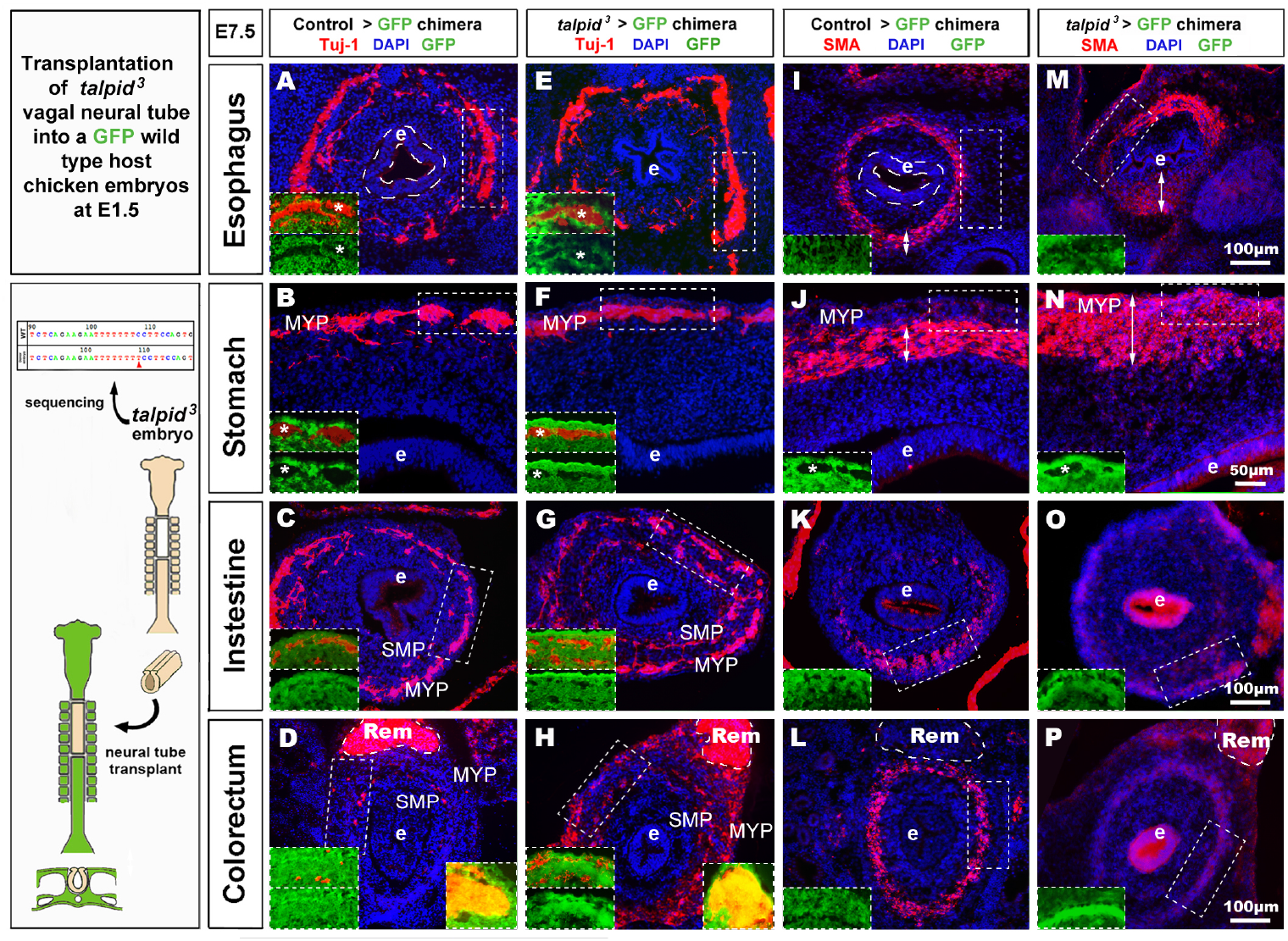
*talpid^3^*>GFP chimeric embryos have grossly normal ENS but smooth muscle and epithelium defects. (A-Q) E7.5 chimeric embryos after neural tube transplantation in (A-D,I-J) control>GFP and (B-H, M-P) *talpid^3^*>GFP. (A-P) Immunofluorescent staining for a TUJ-1+ and GFP [inset] in (A-H) and SMA and GFP [inset] in (I-P). (A-H) normal ENS patterning is observed in both control>GFP and *talpid^3^*>GFP. (D,H insets) show the presence of (red) vagal TUJ-1+/ GFP-ENCC (marked with asterisks *), as well as (yellow) TUJ-1+/GFP+ sacral ENCC in the Remak nerve. (E-H) TUJ-1 staining shows grossly normal ENS in *talpid^3^*>GFP chimera with a more mature / developed appearance of the ENS in the intestine and colon compared to the control chimera. (I-L) Well-defined circular SMA staining in control chimera. (M-P) Diffuse SMA expression and / or extended SMA domain (double headed arrows) in *talpid^3^*>GFP chimera. (O) SMA staining is almost absent in intestine of *talpid^3^*>GFP chimera. (P) Diffuse, yet circular SMA staining present in colorectum of *talpid^3^*>GFP chimera. (A,E,I,M) The endoderm of *talpid^3^*>GFP chimera is significantly thinner than controls at the level of the esophagus. e: endoderm; Rem: nerve of Remak; MYP: myenteric plexus; SMP: submuscosal plexus.

### Hh signaling is disrupted in *talpid^3^* GI tract

Due to the well-established functional connection between *TALPID3* and Hh signalling, we investigated changes in this pathway in control and *talpid^3^* mutants using *in situ* hybridization. *In situ* analysis of *shh* expression in the GI tract of E6.5 control and *talpid^3^* embryos revealed transcripts in the gut epithelium of both type of tissues, demonstrating expression of the *shh* gene in the *talpid^3^* mutants (Fig. 2A-H). Expression of SHH was confirmed in the endoderm of the intestine using an anti-SHH antibody (Supl Fig. S1). To visualise the readout of Hh signalling in the gut wall, mRNA expression of the Hh receptor *PTCH1* was used. In E6.5 control embryos, the pattern of expression of *PTCH1* was composed of one or two concentric gradients surrounding the epithelium (Fig. 2I-L). In the esophagus, the ventral part of the stomach and the intestine, two concentric gradients were present; the first was located adjacent to the epithelium in the sub-epithelial mesenchyme, and the second more distally located in the outer mesenchyme (Fig. 2I-K). In the dorsal part of the stomach and the colorectum a single sub-epithelial gradient was present (Fig. 2J,L). Strikingly, the discrete characteristic gradients of *PTCH1* expression were absent in the GI tract of *talpid^3^* mutants. Instead a diffuse homogenous expression was observed throughout the gut wall and the epithelium. The same diffuse *PTCH1* pattern was observed at all levels of the GI tract (Fig. 2M-P). Together these results showed that, despite epithelial expression of SHH in *talpid^3^* mutants, the precise integration of the signal in the surrounding mesenchyme was lost, as shown by the aberrant *PTCH1* expression.

### Defective expression of extracellular matrix components in *talpid^3^* gut mesenchyme leads to the disappearance of NCC repellent cues

In search of a mechanistic explanation for the lack of ENS plexus formation in *talpid^3^* mutants, we analysed expression of components of the gut ECM known to influence neuronal behaviour (Tennyson et al. 1990; Siebert et al. 2014; Nagy et al. 2016). First, we used the CS-56 antibody, which stains the glycosaminoglycan portion of native chondroitin sulfate proteoglycans (CSPG). CSPG are ECM components regulated by Hh signaling (Nagy et al. 2016). In E6.5 control tissues, CS-56 staining comprised two concentric gradients: the first immediately adjacent to the gut epithelium (sub-epithelial mesenchyme) and the second located in the outer gut wall (outer mesenchyme) (Fig. 3A). This expression pattern was very similar to *PTCH1* as described above. In E6.5 *talpid^3^* gut tissues, CS-56 expression was absent from the outer mesenchyme, with only cells adjacent to the epithelium staining positive (Fig. 3F). Additionally, we examined the expression of Collagen 9 (Coll9), a CSPG expressed in the developing gut, which has been specifically shown to elicit avoidance behaviour by neural crest cells *in vitro* (Ring et al. 1996; Nagy et al. 2016). Consistent with the CS-56 staining, Coll9 pattern of expression was also composed of two concentric areas, one within the sub-epithelial mesenchyme and one in the outer mesenchyme (Fig. 3B,D). Interestingly, the Coll9 expression pattern appeared to define exclusion zones for the migrating ENCC, which were only present outside Coll9 positive areas, as shown by N-cadherin (NCadh) staining (Fig. 3B,C; double arrows). As seen with CS-56, there was no Coll9 expression in the outer mesenchyme of E6.5 *talpid^3^* gut samples and only the sub-epithelial mesenchyme was stained (Fig. 3G,H). Interestingly, and in correlation with the lack of outer mesenchyme Coll9 expression, NCadh-positive ENCC were scattered throughout the gut mesenchyme, with some cells located adjacent to the epithelium (Fig. 3G,H; arrowheads in inset). We also investigated the expression of Coll9 respective to smooth muscle differentiation. We found that SMA and Coll9 have distinct, yet partially overlapping, patterns of expression in control tissues (Fig. 3D,E). In E6.5 control esophagus, the distal gradient of Coll9 corresponded with the inner boundary of the smooth muscle ring (Fig. 3D). In the stomach, Coll9 and SMA were mostly mutually exclusive apart from a subdomain in the ventral region where both were co-expressed (Fig. 3E). In E6.5 *talpid^3^* esophagus, both the Coll9 distal gradient and the SMA ring were absent (Fig. 3I). In the *talpid^3^* stomach, most of the Coll9 expression domain was absent compared to control, while the SMA-positive domain was extended (Fig. 3J). These results suggest that expression of CSPG in the gut mesenchyme is regulated by *talpid^3^*, as its absence led to significant loss in expression of these ECM components. Moreover, the changes in CSPG expression were concurrent with mislocalisation of migrating ENCC.

### Transplantation of wild type ENCC does not rescue the formation of ENS plexuses in a *talpid^3^* gut

To investigate the role of *TALPID3* in ENS plexus formation, we attempted to rescue normal ENS patterning in *talpid^3^* mutants by grafting wild type neural tubes, including neural crest, into *talpid^3^* host embryos (Fig. 4). GFP chicken tissues were used as donors for the transplants, so that vagal ENCC had a functional *TALPID3* protein and could also be traced in the chimeric embryos using GFP expression. When compared to GFP>wild type transplanted controls (Fig. 4A-D) or stage matched *talpid^3^* embryos (Fig. 4I-L), transplanted GFP+ ENCC behaved similarly to ENCC of *talpid^3^* embryos and were unable to rescue ENS patterning (Fig. 4E-H). At the level of the esophagus, instead of surrounding the esophagus in a presumptive circular plexus, as seen in controls (Fig. 4A), transplanted GFP+ ENCC clustered in the dorsal region (Fig. 4E), in a pattern similar to that seen in *talpid^3^* embryos (Fig. 4I). In the stomach and intestine, transplanted GFP+ ENCC migrated similar distances along the GI tract as observed in *talpid^3^* embryos but were scattered throughout the mesenchyme (Fig. 4F,G,J + inset G) and not arranged in plexuses, as seen in controls (Fig. 4B-D + inset C). Epithelium and smooth muscle thickness (stained with DAPI and Phalloidin, respectively) in intestine and stomach sections were measured in stage matched GFP>wild type and GFP>*talpid^3^* chimera, as well as *talpid^3^* mutants. Measurements from GFP>*talpid^3^* chimeric tissues and *talpid^3^* mutant were both statistically (\*\*\**p<0*.*001*) different from the controls (Supl. Fig. S2). The non-parametric Spearman’s Rho test was used to measure the correlation between GFP>wild type and GFP>*talpid^3^* chimera, and *talpid^3^* mutants measurements. The correlation between GFP>wild type and GFP>*talpid^3^* transplants was 0.670, whereas the correlation between the GFP>wild type transplant and the *talpid^3^* mutant was 0.715. Importantly, the correlation between the GFP>*talpid^3^* and the *talpid^3^* mutant was 0.942, an extremely high score. The fact that measurements from chimera and mutant clustered together and were statistically different from the control supported histological findings of a lack of rescue. Overall, wild type transplanted vagal ENCC did not rescue ENS plexus formation or the smooth muscle phenotype in a *talpid^3^* environment.

### Lack of TALPID3 in ENCC does not affect gross ENS morphology but affects smooth muscle and mucosa development in a non cell-autonomous manner

To further assess the role of TALPID3 during ENS development and to test its cell autonomous requirement in ENCC, we performed the converse experiment to that described above, by grafting *talpid^3^* vagal ENCC into GFP-expressing chick embryos. In the resulting chimeric embryos, the transplanted ENCC were devoid of functional TALPID3 protein and were identified by lack of GFP expression (GFP^-ve^; Fig. 5 inset). Due to the extremely high mortality of this type of chimera, only one specimen survived to E7.5 for the gut tissues to be analysed. Surprisingly, transplanted vagal *talpid^3^* ENCC migrated throughout the GI tract and patterned normally into the characteristic MYP and SMP of the ENS, as shown by HNK1 and GFP staining at E7.5 (Fig. 5A-H). Importantly, *talpid^3^*ENCC colonised the entire length of the GI tract and were found in the colorectal region (Fig. 5H + inset). Here, vagal GFP^-ve^ ENCC (red) were present alongside GFP^+^ (yellow) ENCC (Fig. 6N inset) that were likely sacral ENCC entering the gut prematurely (Burns and Le Douarin 2001). In contrast to the ENS, which was grossly normal in the chimeric embryo, there were obvious defects in smooth muscle differentiation as revealed by SMA staining: instead of the characteristic SMA^+^ ring of differentiating smooth muscle cells seen at E7.5 in control chimeras (Fig. 5I-P + insets J,N), there was thickened and diffuse SMA staining in regions of the esophagus and the stomach, suggesting impaired differentiation (Fig. 5I,J,M,N double arrows + insets J,N; Fig. 6H). The characteristic SMA^+^ ring of cells was also missing in the intestine of the chimeric embryo (Fig. 5O). Normal SMA staining was observed in some areas around the esophagus and within the colorectum, which coincided with the presence of normal vagal ENCC from the region adjacent to the graft and wild type sacral ENCC, respectively (Fig. 5H inset). Additionally, the architecture of the mucosa was disrupted in the chimeric embryo, with the esophageal epithelium consisting of a thin folded monolayer, unlike controls where a thickened circular epithelium was present (Fig. 5A,E,I,M doted lines). Using DAPI, SMA and TuJ-1 staining, epithelial cells, smooth muscle cells and enteric neurons were counted on stomach sections (n=3 sections minimum, as shown in Fig. 5 B,F,J,N). Numbers were combined with equivalent murine data for chicken/mouse cross-species statistical analysis of chimeric animals where *Talpid3* was knocked down in ENCC, as described below (Fig. 6H).

**Figure 6:**
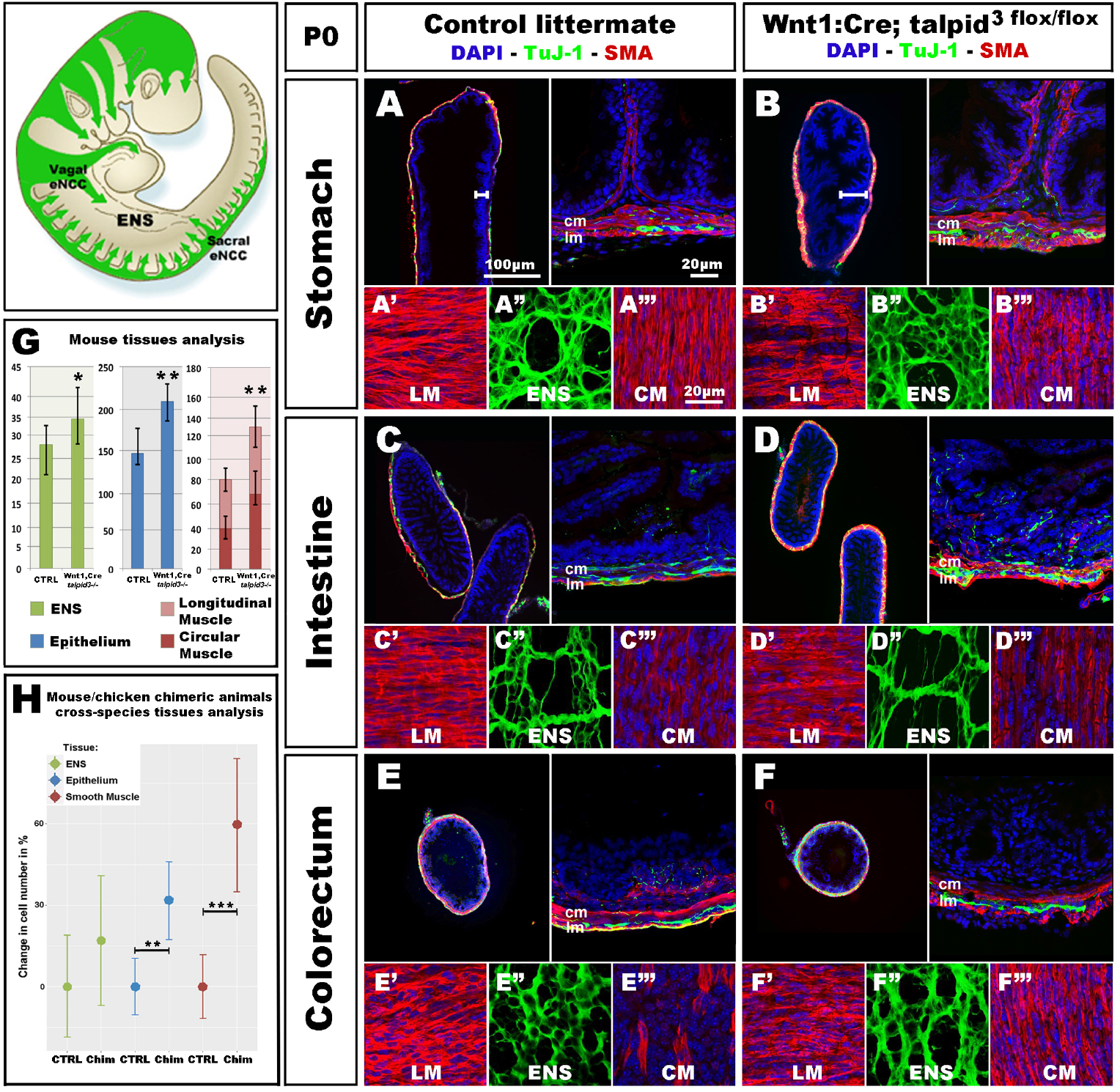
*Wnt1:Cre;Talpid3* ^*flox/flox*^ mice show grossly normal ENS but smooth muscle and epithelium defects. (A-F) Immunofluorescent staining for TuJ-1 and SMA at the level of (A,B) the stomach, (C-D) intestine and (E,F) colorectum in control littermate and *Wnt1:Cre;Talpid3* ^*flox/flox*^ P0 pups respectively. (A’-A’’’, B’-B’”, C’-C”‘) confocal projections at the level of the longitudinal muscle (LM), enteric nervous system (ENS) and circular muscle (CM) respectively. (G) Bar graph shows cell counts for ENS, LM, CM and total smooth muscle, as well as epithelium in control littermate versus *Wnt1:Cre;Talpid3* ^*flox/flox*^. In the stomach, the ENS (green) showed a statistically significant (\**p=0*.*045*) increase of 29% in the number of ENCC in the mutant, compared to control. Epithelium (blue) showed a statistically significant (\*\**p=0*.*009*) increase of 39% in the mutant, compared to control. Muscle layers (red) showed a highly statistically significant (\*\**p=0*.*002*) increase of 67% in mutant tissue sections compared to control, with 80% increase in the longitudinal muscle (\*\**p=0*.*002*) and 55% increase in the circular muscle (\*\**p=0*.*018*). Difference in epithelium thickness is highlighted in A and B by white bars. (H) Scatter plot shows mouse/chicken cross-species cell counts for ENS, epithelium and smooth muscle in stomach of controls and chimeric animals. Epithelium (blue) showed a statistically significant (\*\**p=0*.*009*) increase of 39% in the mutant, compared to control. Muscle layers (red) showed a highly statistically significant (\*\**p=0*.*002*) increase of 67% in mutant tissue sections compared to control.

Because only one *talpid^3^*>GFP chicken chimera could be analysed (due to extremely high mortality) we wished to confirm the chicken results using an alternative approach. For this, we engineered a conditional knock out of the *Talpid3* gene in a mouse model by crossing a *Wnt1:Cre* line (Danielian et al. 1998), with a floxed *Talpid3* knock-out line. Resultant *Wnt1:Cre;Talpid3* ^*flox/flox*^ embryos, which do not express TALPID3 in NCCs, were often embryonic lethal or died at P0, due to presumptive respiratory problems, as their lungs failed to fill with air (Sulp. Fig. S4). P0 pups showed a craniofacial phenotype characteristic of hypoplastic neural crest derivatives (Sulp. Fig. S4D,F). In mutants, the GI tract morphology was grossly normal compared to control littermates (Sulp. Fig. S4B,E). As seen in the chicken model, *Talpid3* ENCC colonised the entire length of the gut and formed an apparently normal ENS all along the GI tract (Fig. 6A-F; A”-F”). Although the ENS was grossly normal, quantitative analysis of the ENS in the stomach region showed a statistically significant (*p= 0*.*045*) increase of 29% in the number of ENCC in the mutants (n= 8 sections) (Fig. 6G). Of note, and again in accordance with the defects observed in the chicken model, the number of cells in both the epithelium and muscle layers was affected, in a non-cell autonomous manner, by the lack of TALPID3 in the ENCC. Indeed, quantitative analysis of cell numbers in both tissues showed statistically significant 39% increase in the epithelium (*p=0*.*009*; n= 7 sections) and 67% increase in the muscle layers (*p=0*.*002;* n= 7 sections), with an 80% increase in the longitudinal muscle (*p=0*.*002)* and 55% increase in the circular muscle (*p=0*.*018*). Interestingly, the smooth muscle myoblasts appeared misshaped in the conditional *Talpid3* mutant, pointing towards impaired muscle differentiation (Fig. 6A’-F’;A”-F” and S1;S2 videos). The transgenic mouse measurements were combined with equivalent chicken data from the *talpid^3^*>GFP transplant for mouse/chicken cross-species analysis. Cell counts for epithelium showed a statistically significant (\*\**p=0*.*009*) increase of 39% in the chimeric animals compared to controls. Smooth muscle cell count showed a highly statistically significant (\*\**p=0*.*002*) increase of 67% in chimeric animals compared to control. The relative increase observed in ENS cell numbers was not statistically significant in this cross-species analysis (Fig. 6H). Overall, apart from altered cell numbers, knocking out *talpid^3^* in ENCC had little effect on early ENS development and patterning in both chicken and mouse models, but it altered growth and differentiation of smooth muscle & mucosa in a non cell-autonomous manner.

### Human embryonic tissues bearing a KIAA0586 mutation recapitulate the GI defects observed in TALPID3 animal models

To assess whether the role of TALPID3 is conserved throughout evolution and is relevant to human GI tract development and patterning, we examined human fetal GI tissues obtained from a 26 week human fetus with a homozygous for a 1815G>A mutation in *KIAA0586*, the human orthologue of *talpid^3^* *(chicken) and Talpid3 (mouse)*, as previously described (Alby et al. 2015). Previous anatomical description of the fetus listed shortened ribs, micromelia, lingual hamartomas, postaxial and preaxial polydactyly, temporal polymicrogyria and an occipital keyhole defect (Alby et al. 2015; Cocciadiferro et al. 2020). Gross examination of the GI tract revealed an elongated and tubular stomach, but otherwise apparently normal GI tract. Histological analysis showed a portion of the intestine (“segment 1”) with grossly normal neuromuscular pattern (Fig. 7A,D,G) consistent with that observed in a 26 week control fetus (Fig. 7C). However, another portion (designated as “segment 2”) showed massive overgrowth of the smooth muscle layers (∼140 mm in the mispatterned “segment 2”, compared to ∼20 mm in “segment 1”and 26 week control) and the mucosa (∼25 mm in the mispatterned “segment 2”, compared to ∼6 mm in “segment 1” and 26 week control). The smooth muscle overgrowth is “segment 2” corresponded to an increase in thickness of 6.5 times (Fig. 7B,E,H). Additionally, immunostaining for TuJ1 showed enteric neurons scattered throughout the gut wall (Fig. 7E), in a pattern similar to that observed in the *talpid^3^* chicken model (Fig. 1O-R). The striking similarities found in histological observations were confirmed by combining equivalent measurements from human and chicken intestinal tissues for human/chicken cross-species analysis. This analysis showed a statistically significant increase in (i) ENS cell numbers (40.8%,\*\**p=0*.*004*) and (ii) epithelium and smooth muscle tissue thickness (77%,\*\*\**p=0*.*00006* and 413%, \*\*\*\**p<<0*.*0001* respectively) in mutant tissues compared to stage matched controls (Fig. 7I). We also investigated expression of human chondroitin sulfate proteoglycans using CS56 (Fig. 7F-H).19 and 26 week controls showed expression of CS56 in the serosa the submucosa and the mucosa (Fig. 7F; Supl. Fig. S5B). In “segment 1”, CS56 expression was also observed in the serosa and the mucosa whereas, strikingly, in “segment 2” no CS56 expression was detected (Fig. 7G,H). To investigate further the intestinal phenotype of this fetus, we examined the expression of both Patched and Cytokeratin (Fig. 7J-O). Even though cytokeratin was expressed in the fetus bearing the *KIAA0586* mutation, the staining was not as strong and uniform as in the control. Likewise, Patched staining was reduced in “segment 1” of the fetus bearing the *KIAA0586* mutation compared to the control, whereas “segment 2” showed no staining. The altered *patched* expression was confirmed by *in situ* hybridization (Sulp. Fig. 4C-E). Both epithelial stainings pointed to delayed / altered gut epithelial differentiation. Overall, the phenotype observed in “segment 2” of the fetus was strikingly similar to that of the *talpid^3^* chicken model with scattered enteric neurons, smooth muscle & mucosa overgrowth as well as impaired differentiation. Other defects observed in both human and the chicken model included: (i) altered Hh pathway (as shown by a lack of Patched expression and loss of mRNA expression pattern) (ii) impaired epithelium differentiation (as shown by Cytokeratin) and (iii) lack of CSPG components of the ECM.

**Figure 7:**
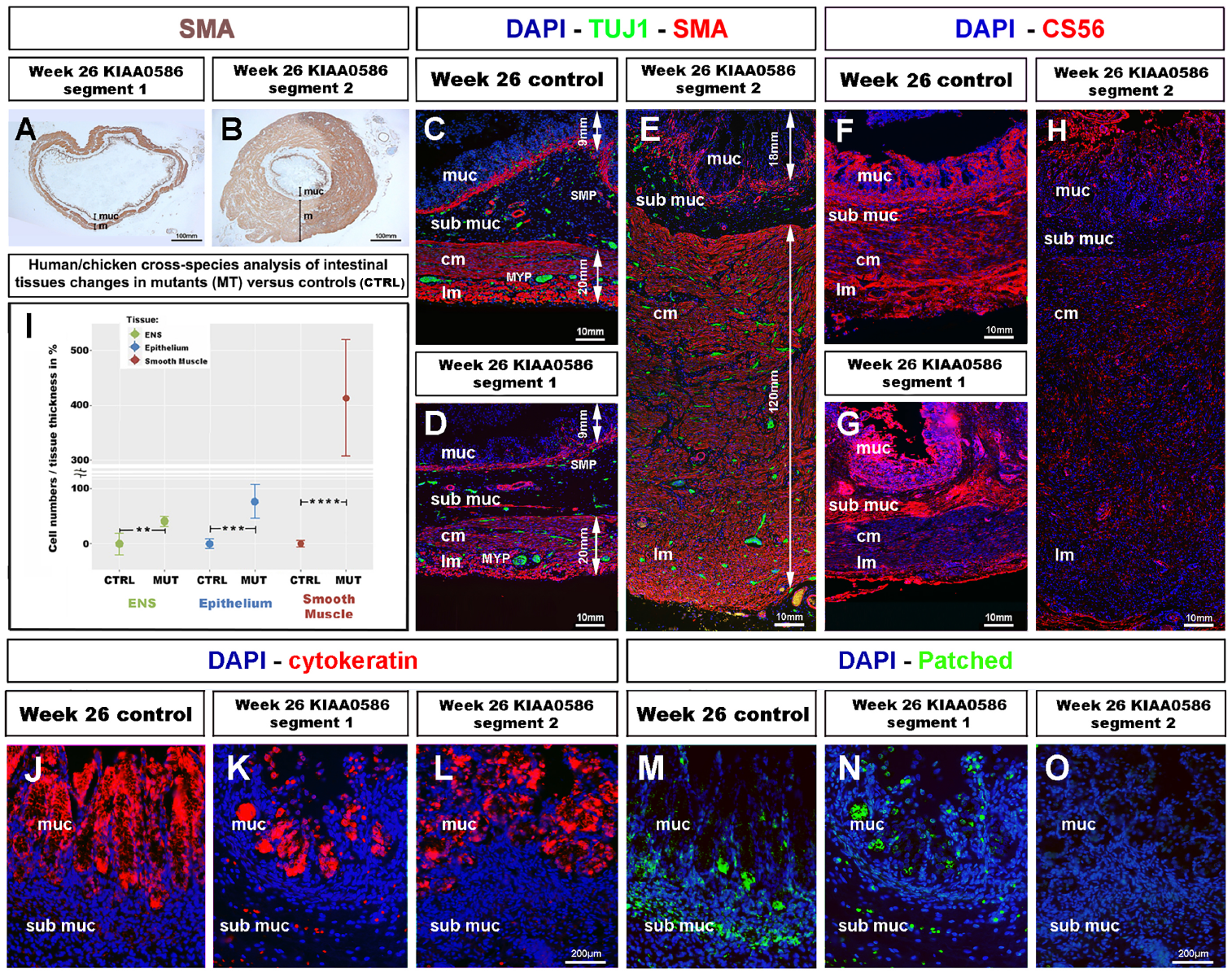
Fetal GI tissues from a 26 week human fetus bearing a *KIAA0586* mutation show one gut segment with scattered ENS, mucosa & smooth muscle overgrowth, as well as CS56, Cytokeratin & Patched expression defects. (A,B) Low magnification gut sections of a 26 week fetus with a *KIAA0596* mutation. SMA immuno-staining reveals grossly normal smooth muscle layers in (A) “segment 1” of the intestine and extensive smooth muscle and mucosa overgrowth in (B) “segment 2”. (C-E) Immunofluorescence staining for DAPI (nucleus), TuJ-1 (ENS) and SMA (Smooth Muscle) on intestine sections. (C) 26 week control intestine with normal ENS patterning and longitudinal and circular smooth muscle layers. (D,E) staining in the *KIAA0586* fetus tissue shows grossly normal neuromuscular pattern in (D) “segment 1”, with well-defined ENS plexuses, normal muscle (∼20mm) and mucosa (∼6 mm) thickness. (E) “segment 2” has scattered ENS with smooth muscle (∼140 mm) and mucosa (∼25 mm) overgrowth. (F-H) Immunofluorescence staining for DAPI (nucleus) and CS56 (chondroitin sulfate proteoglycans) on intestine sections. (F,G) 26 week control intestine and Segment 1” with CS56 expression in the serosa, submucosa and mucosa. (H) “Segment 2” shows absence of CS56 staining. (I) Human/chicken cross-species analysis of intestinal tissues in *talpid^3^* and KIAA0586 mutants has statistically significant increase in (i) ENS cells numbers (40.8%,\*\**p=0*.*004*) and (ii) epithelium and smooth muscle tissue thickness (77%,\*\*\**p=0*.*00006* and 413%, \*\*\*\**p<<0*.*0001* respectively). (J-K) Immunofluorescence staining for DAPI (nucleus) and Cytokeratin on mucosal sections. (J) 26 week control intestine has uniform Cytokeratin staining in the mucosa. (K-L) “Segment 1 and 2” show uneven Cytokeratin expression. (M-O) Immunofluorescence staining for DAPI (nucleus) and Patched. (M) “Segment 1” has reduced submuscosal staining and normal mucosal staining compared to 26 week control. (O) “Segment 2” shows no Patched staining. muc: mucosa; sub muc: submucosa; cm: circular muscle; lm: longitudinal muscle; MYP: myenteric plexus; SMP: submuscosal plexus.

## DISCUSSION

Although TALPID3 has been recognized to have essential functions during embryonic development, its role during GI and ENS development has not, as yet, been studied. Previous investigations have shown that TALPID3 animals are useful to model human birth defects such as short ribs, polydactyly, or craniofacial abnormalities that are attributed to abnormal hedgehog signaling (Davey et al. 2007; Bangs et al. 2011; Ben et al. 2011; Alby et al. 2015). Our findings establish that TALPID3 animal models can also offer insights for congenital human GI defects, such as short gut, tracheoesophageal atresia/fistula, and anorectal abnormalities, i.e. a variety of defects commonly seen in paediatric gastroenterology clinics. Moreover, the striking similarities between the neuromuscular abnormalities described here, in chicken, mouse and human, demonstrate that the function of TALPID3 is well conserved across species and is of importance for normal human gut development.

### TALPID3 is a regulator of GI neuromuscular patterning

We specifically examined the role of TALPID3 in neuromuscular patterning of the developing GI tract, as it was clear from the histological defects in both chicken and human GI tissues that lack of TALPID3 led to severe disruption of this developmental process. Our analysis showed that the lack of TALPID3 consistently affected the neuromuscular patterning of the gut. Interestingly though, the resulting phenotypes varied, depending on the location along the A-P axis of the gut. In the chicken model, lack of TALPID3 in the esophagus led to complete absence of smooth muscle and ENS, whereas in the stomach and the intestine lack of TALPID3 led to ectopic smooth muscle differentiation and misplaced enteric neurons. Likewise, one portion of the human intestine from a fetus bearing a *KIAA0586* mutation showed grossly normal neuromuscular patterning, whereas another had dramatic muscle overgrowth and scattered enteric neurons. These observation, when combined with quantifications of enteric neuronal, smooth mucslce and epithelial cell numbers, suggest that, albeit with species differences, TALPID3 is part of a conserved mechanism controlling the appropriate spatial differentiation of smooth muscle and the correct positioning of enteric neurons. Moreover, TALPID3 functions as part of regional-specific mechanisms regulating the correct neuromuscular patterning at different levels of the GI tract. Perhaps this is not surprising since the spatiotemporal development of GI smooth muscle has been shown to have regional-specific differences (Bourret et al. 2017; Graham et al. 2017). Our results are in accordance with recent findings in the related chicken mutant *talpid2* (Brooks et al. 2021).

### Disruption of GI patterning in *talpid^3^* mutant is linked to impairment of the Hh pathway

Our findings add to the body of evidence linking TALPID3 to the Hh signalling pathway (Davey et al. 2007; Bangs et al. 2011; Ben et al. 2011; Alby et al. 2015; Li et al. 2017). The link between TALPID3 and Hh signalling in the gut can be observed in (i) the gross anatomy of the GI tract (ii) the smooth muscle and (iii) ENS defects of the mutants. (i) First, the gross anatomical abnormalities in the TALPID3 chicken GI tract are consistent with a loss of Hh signalling. Severe reduction in GI tract size with normal gross anatomy has been reported using a conditional approach to remove both *Shh* and *Ihh* functions from early mouse gut endoderm (Mao et al. 2010). Likewise, mouse knockouts of *Shh, Gli2* and *Gli3* all display tracheo-esophageal atresia/fistula, a phenotype also described here in the TALPID3 chicken mutant (Motoyama et al. 1998; Ramalho-Santos et al. 2000). Additionally, regulation of gut epithelium homeostasis has been linked to the Shh signalling pathway and both *talpid^3^* chicken and *KIAA0585* human gut tissue showed altered epithelium growth / differentiation (Mao et al. 2010; Ben-Shahar et al. 2019). Lastly, Hedgehog signalling is critical for normal anorectal development and anorectal malformations have been reported in *shh* knockout mice and in humans with polymorphisms in Hedgehog genes (Ramalho-Santos et al. 2000; Gao et al. 2016). The open hindgut phenotype of the *talpid^3^* chicken model offers an additional model for researching such malformations. (ii) It has been shown that Shh signalling regulates the concentric architecture of the intestine. In particular, SHH affects the mesenchyme immediately adjacent to the epithelium, where it restricts smooth muscle differentiation to allow the connective tissue of the submucosa to develop (van den Brink 2007; Huycke et al. 2019). The increased domain of smooth muscle differentiation in the stomach and intestine of the *talpid^3^* chicken suggests that this Hh-dependant regulation is lost. Moreover, the results from *ptch in situ* hybridization in *talpid^3^* mutants directly demonstrate that SHH expressed by the endoderm is not correctly integrated throughout the radially surrounding mesenchyme. The lack of PATCHED protein immunostaining in “segment 2” suggests that this part of the gut is unable to respond to SHH signalling. The loss of the proximal SHH readout in the *talpid^3^* mutants can be directly correlated with the subsequent differentiation of smooth muscle adjacent to the epithelium in the stomach and intestine. This result demonstrates that mesenchymal cells surrounding the epithelium are competent for induction into smooth muscle, but are normally prevented from doing so by the proximal SHH gradient *via* a TALPID3-dependent mechanism. In the esophagus, the lack of TALPID3 does not lead to an extension of the smooth muscle differentiation domain, but rather to a lack of smooth muscle differentiation altogether, pointing to a different region-specific readout of the Hh signal. This failure of smooth muscle differentiation in the esophagus may explain why migrating ENCC do not halt their migration and differentiate into enteric neurons, as they lack target tissue to innervate. Indeed, ENCC were observed at the level of the esophagus at stage E6.5 in the *talpid^3^* chicken mutant, but no cells were present in this region at E8.5. TALPID3 is therefore necessary for the correct integration of SHH throughout the gut mesenchyme and to define the concentric architecture of the smooth muscle differentiation domain. Our results fit in with a recent study showing that Hedgehog acts through Bmp signaling to inhibit subepithelial smooth muscle and that levels of Hedgehog signaling regulate differentiation of the inner smooth muscle layer (Huycke et al. 2019). This suggest that TALPID3 is part of this regulatory mechanism. (iii) The influence of SHH on NCC migration and proliferation is well established, but many paradoxical results point to a complex role for Hh during ENS development with species and developmental differences, as well as direct and indirect regulatory mechanisms (Ramalho-Santos et al. 2000; Fu et al. 2004; Reichenbach et al. 2008; Biau et al. 2013; Jin et al. 2015). Recent results clearly show, both in chicken and mouse, that the SHH receptor Patched is not expressed by ENCC and point to indirect regulatory mechanisms (Nagy et al. 2016). Our findings are in agreement with an indirect regulation of ENCC by SHH through its role in modifying the gut ECM and thereby the environment through which ENCC migrate, as we discuss below.

### Loss of TALPID3 and Hh signalling leads to ECM defects and absence of environmental NCC repellent cues

In search of a mechanistic explanation for the lack of ENS plexus formation in *talpid^3^* mutants, we analysed expression of CGPG molecules, which are components of the gut ECM. Expression of ECM components is known to be regulated by both primary cilia and Hh signalling (Seeger-Nukpezah and Golemis 2012; Nagy et al. 2016), making them good candidates for further investigation. Alterations in ECM expression have also been described in human ciliopathies (Ramalho-Santos et al. 2000; Seeger-Nukpezah and Golemis 2012). Importantly, CSPG molecules have been shown to provide guidance cues for neuronal behaviours such as migration, axon outgrowth and axon termination (Ring et al. 1996; Carulli et al. 2005; Siebert et al. 2014). We specifically investigated Coll9, a CSPG expressed in the developing gut, which has been specifically shown to elicit avoidance behaviour by NCC *in vitro* (Ring et al. 1996; Nagy et al. 2016). In correlation with the loss of Hh signalling in the GI tract of both the *talpid^3^* chicken and the fetus bearing a *KIAA0586* mutation, expression of CSPG molecules was lost. Importantly, double staining of CSPG and NCC showed that the disappearance of the CSPG expression correlated tightly with ectopic localization of ENCC, suggesting that the lack of repellent molecules such as Coll9 is a direct mechanism underlying the lack of ENS plexus formation in *talpid^3^* GI tract. This finding is in agreement with previous studies showing that the regulation of CSPG by SHH in both chicken and mouse models can modify the behaviour of ENCC and subsequently ENS patterning (Nagy et al. 2016). To investigate a possible connection between smooth muscle differentiation and the expression of CSPG, we performed double immunofluorescence with SMA and coll9 antibodies. We found that SMA and Coll9 have distinct, yet partially overlapping, patterns of expression. This demonstrates that some myoblasts express Coll9. However, Coll9 is mainly expressed in mesenchymal cells and it is this wider CSPG expression domain, and not the SMA domain, that correlates best with the regional localisation of ENCC. Additionally, our chicken transplantation experiments demonstrated the inability of wild type ENCC to rescue ENS formation in a TALPID3-null gut environment. This finding highlights the role of environmental cues in directing ENS formation and shows that ENS plexus development does not rely on self-organising properties of ENCC. This is in agreement with recent work showing that disruption of the ENCC environment can disrupt ENS patterning (Graham et al. 2017). Our study also demonstrates that TALPID3 expression in the gut environment is essential for proper expression of guidance cues that direct ENS plexus patterning. Importantly, we also show that this role is conserved during human fetal gut development.

### TALPID3 is not required cell autonomously for ENS plexus formation but is a regulator of neuronal-mesenchymal-epithelial interactions directing correct tissue growth and differentiation

We investigated the cell autonomous requirement for TALPID3 during ENCC migration and gut patterning by knocking out *talpid^3^* specifically in ENCC, either by neural tube transplantation in chicken or using a *Wnt1-Cre* driven *talpid^3^* conditional knockout in mouse. In both models, TALPID3 was found not to be required for ENCC migration or normal ENS plexus formation as the gross morphology of the ENS was unaffected. The only alteration to the ENS in these conditions was the increase in cell number observed in the chicken model and in the mouse model, which we quantified in stomach sections. Alteration of NCC numbers could be the result of subtle migration or differentiation defects changing the relative number of ENCC in specific locations. Alternatively, it could be linked to a TALPID3-dependent alteration of the SHH pathway and, specifically, the influence of SHH on NCC as a mitogen (Fu et al. 2004; Reichenbach et al. 2008; Roper et al. 2009; Nagy et al. 2016). Remarkably, knocking out *talpid^3^* in ENCC led to wider non-cell autonomous defects in other tissues such as smooth muscle and epithelium. Both the mouse and chicken phenotypes revealed a neural crest TALPID3-dependent mechanism controlling growth and differentiation of mesenchyme and epithelium. The most striking effect was alteration of smooth muscle differentiation and increased numbers of myoblasts which we quantified in stomach sections in chicken and mouse. The alteration of smooth muscle differentiation was particularly evident in the stomach region of mice. Variations in epithelium shape and cell numbers were also evident both in mouse, chicken (. Mouse/chicken cross-species analysis confirmed that these differences in smooth muscle and epithelial cell numbers were statistically significant when comparing chimeric animals to controls. Interestingly, in addition to its role in smooth muscle development, the Hh pathway also plays an important role in mucosal growth (Mao et al. 2010; Ben-Shahar et al. 2019). Indeed *shh* and *gli3* mutant mice have mucosal hyperplasia in the stomach (Ramalho-Santos et al. 2000; Kim et al. 2005). It has been previously postulated that vagal ENCC could act as a mediator in the mesenchymal-epithelial interactions that control stomach development (Faure et al. 2015). Additionally it has been shown that ENS influences the differentiation and growth of other cell types within the gut, as demonstrated by smooth muscle overgrowth in the aganglionic portion of EdnrB mice, as well as alteration of goblet cell differentiation (Thiagarajah et al. 2014). Recent work has also shown that implantation of neural crest cells within tissue-engineered small intestine altered the transcriptome of a wide variety of gastrointestinal cell types, demonstrating the necessity of the neuronal lineage to be able to recapitulate gut organogenesis *in vitro* (Schlieve et al. 2017). Our findings emphasise the importance of the neural component for correct smooth muscle and mucosa development both during embryonic gut development and for regenerative medicine research. Moreover our study shows the central role played by TALPID3 in neuronal-mesenchymal-epithelial interactions necessary for normal GI tract development.

The neural crest specific *talpid^3^* knockout results described here are in agreement with non-cell autonomous effects seen in other tissues after NCC-specific gene knockout. For example, knocking out the intraflagellar protein Kif3a in NCC led to non-cell autonomous striated muscle defects during tongue development (Millington et al. 2017). Additionally, loss of NCC disrupts the distribution of second heart field cells in the pharyngeal and outflow regions (Bradshaw et al. 2009). Likewise, conditional neural crest Rac1 knockout, using a *Rac1/Wnt1-Cre* line, shows excessive proliferation of SMA+ cell wall around the aortic sac and ventral aorta (Waldo et al. 2005; Thomas et al. 2010). Investigating the gut phenotype of *Rac1/Wnt1-Cre* mutants for smooth muscle defects would be informative. Interestingly, Rac1 is downstream of the non-canonical SHH pathway (Mulligan 2014). TALPID3 and Rac1 have a comparable function in vesicular trafficking (Stenmark 2009), centrosome regulation (May et al. 2014) and in cell-matrix interactions (Thomas et al. 2010). Rac1 is also downstream of RET, a tyrosine kinase receptor essential for ENS development (Natarajan et al. 2002; Fu et al. 2010; Mulligan 2014).

### Migrating ENCC do not extend a prominent primary cilia

Having established that TALPID3 was not required cell autonomously for ENCC migration and gross ENS patterning, and considering the role of TALPID3 in ciliogenesis, it follows that primary cilia might not be required for ENCC migration. ENCC are a highly proliferative cell population and it is their mitogenic activity, which is driving their migration and invasiveness (Simpson et al. 2007). It is also known that there is a reciprocal regulation of cilia and the cell cycle, so that the primary cilium is dismantled in replicating cells, which makes it unlikely for highly proliferative cells to extend a primary cilium (Ishikawa and Marshall 2011; Ford et al. 2018). Interestingly also, migrating interneurons within the developing murine brain do not show an extended primary cilium (Higginbotham et al. 2012). In our study, even though primary cilia were evident in chicken neural tube cells cultured *in vitro*, we failed to observe primary cilia in migrating vagal ENCC, either anatomically by SEM or molecularly using immunofluorescence, arguing against the presence of a primary cilium on migrating ENCC. *In vivo*, despite primary cilia readily being detected in E6.5 chicken gut sections, only 11% of HuC^+^ ENCC showed a detectable primary cilium. It has been shown that differentiated ENS neurons bear a primary cilium (Junquera Escribano et al. 2011; Luesma et al. 2013). Other studies describe primary cilia on YFP+ cranial NCC using the *Wnt1:Cre; Rosa:YFP* transgenic line (Brugmann et al. 2010; Millington et al. 2017). In this transgenic line, YFP is expressed in both the neural tube (which is ciliated) and the neural crest derivatives. This lack of discrimination is problematic to decipher this issue (Cassiman et al. 2006; Murdoch and Copp 2010). Our study suggests that extending primary cilia mainly happens as ENCC start to differentiate into enteric neurons.

### Role of the human TALPID3 orthologue KIAA0586 in human GI tract development

*KIAA0586* is the human orthologue of *talpid^3^*. Some homozygous mutations of *KIAA0586* have been shown to be embryonic lethal and to lead to severe developmental abnormalities such as Hydrolethalus syndrome and short-rib polydactyly (Alby et al. 2015). Here we show that the phenotypic spectrum of *KIAA0586* mutations extends to defects in the GI tract. Apart from a tubular stomach the gross anatomy of the GI tract of the fetus was normal. However, our histological analysis showed severe alteration of the gut patterning with smooth muscle and mucosa hyperplasia, as well as scattered enteric neurons in a sub-section of the intestine we designated “segment 2”. In accordance with the animal models, we show that this phenotype is the consequence of disruption of the Hh pathway and loss of normal ECM expression. Our findings shed light on the central role of KIAA0586 in patterning of the gut during human fetal development. Recently *KIAA0586* mutations have also been identified in Joubert syndrome patients. JBTS is defined by three primary findings: (i) underdevelopment of the brain cerebellar vermis, brain stem defects giving the appearance of the molar tooth sign (MTS) and (iii) Hypotonia (Abdelhamed et al. 2013; Sanders et al. 2015; Stephen et al. 2015). Importantly also, patients typically have a perturbed respiratory pattern in the neonatal period and severe psychomotor delay. Although very rare, association with Hirschsprung disease and problems with bladder and bowel control (incontinence) have been reported in Joubert patients with milder forms of the disease (Shian et al. 1993; Ozyurek et al. 2008; Akhondian et al. 2013; Purkait et al. 2015). Considering the role of KIAA0586 in tissue patterning our study unfolds, it is also tempting to speculate that other symptoms of JBTS (like psychomotor delay, hypotonia and respiratory difficulties), could be linked to neuronal-mesenchymal-epithelial patterning defects in other parts of the body. Interestingly, in this study, *Wnt1:Cre; Talpid3 fl/fl* P0 mutants died at birth from presumed respiratory failure, as their lungs did not appear to inflate. This points to a possible failure in the developmental interactions of neural crest, smooth muscle and epithelium, as lung neural crest cells share developmental origins with ENCC (Burns and Delalande 2005; Freem et al. 2012). Finally, the ENS is often referred to as the “second brain” and it is possible that brain ECM defects, similar to the ones we describe in this study, could underlie abnormal brain patterning defects, such as cortical heterotopias, commonly seen in Joubert syndrome, but also in Bardet– Biedl syndrome or Meckel–Gruber syndrome (Willaredt et al. 2008; Abdelhamed et al. 2013; Higginbotham et al. 2013). In mice, a large variety of CSPGs represent major components of the ECM in the brain (Horii-Hayashi et al. 2015). In human, defects in ECM, leading to impaired neuronal guidance, could underlie other types of brain function abnormalities. A recent CNS conditional *talpid^3^* knockout mouse model for JBTS showed cerebellar defects due to granule cell proliferation, migration and differentiation, three aspects of cellular behaviour also affected in our models (Bashford and Subramanian 2019). This study, however, did not investigate ECM components.

In conclusion, our findings demonstrate a central role for the centrosomal protein TALPID3 in neuromuscular patterning of the developing gastrointestinal tract (summarised in Fig. 8). We show that the function of TALPID3 during GI neuromuscular patterning is conserved in vertebrates including during human fetal development. Our findings also reveal a new role for the ENS in regulating neuronal-mesenchymal-epithelial interactions necessary for normal GI tract development, and that this regulatory mechanism is TALPID3-dependent. These findings add new understandings to human GI tract development mechanisms and have direct implications for regenerative medicine, as they emphasise the importance of the neural component for *in vitro* gut organogenesis.

**Figure 8:**
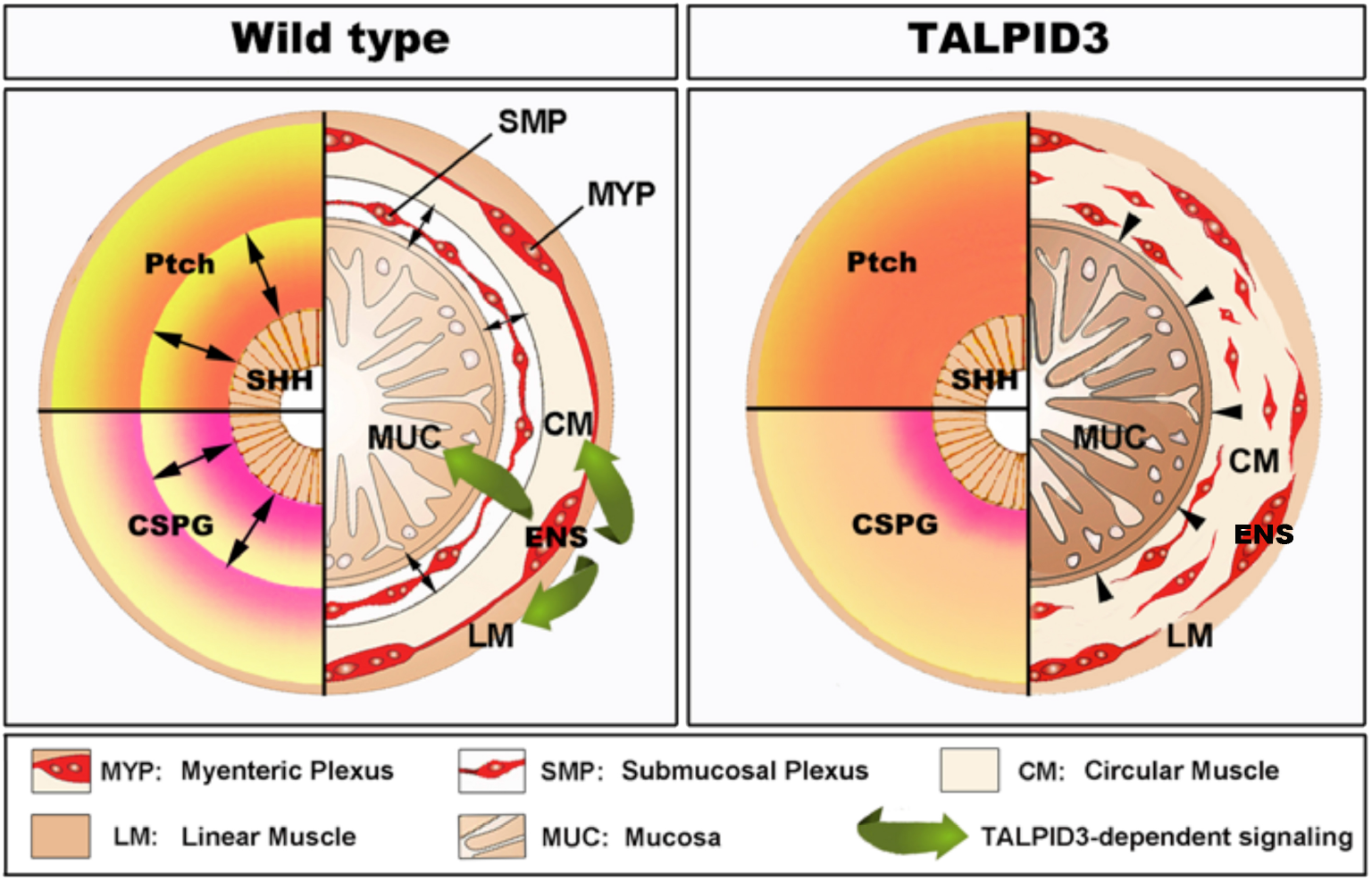
Model of TALPID3-dependent gastrointestinal neuromuscular patterning. During normal gut development (Left), the Sonic Hedgehog (SHH) gradient defines the concentric architecture of the intestine as seen by the discrete PATCHED (Ptch) pattern of expression. SHH readout in the mesenchyme immediately adjacent to the epithelium restricts smooth muscle differentiation to allow the connective tissue of the submucosa to develop. In the absence of a functional TALPID3 protein (Right), the SHH readout is lost, as shown by the diffused *Ptch* expression. Loss of the SHH readout leads to ectopic spread of the smooth muscle differentiation domain adjacent to the epithelium. (Left-bottom) During normal gut development, the precise readout of the SHH gradient directs the pattern of CSPG expression, restricting ENCC to the locations where the enteric nervous system (ENS) will form. The ENS regulates smooth muscle & mucosa growth and differentiation *via* a TALPID3-dependent signalling (green arrows). (Right - bottom) In the absence of a functional TALPID3 protein, SHH readout is lost, which coincides with a loss of CSPG expression. This loss of expression permits ectopic ENCC migration resulting in scattered ENCC throughout the gut mesenchyme. Loss of TALPID3 also leads to altered growth and differentiation of smooth muscle and mucosal epithelium.

## MATERIALS and METHODS

### Chicken intraspecies neural tube grafting

Fertile chicken eggs were obtained from commercial sources within the UK. *talpid^3^* and transgenic GFP chicken eggs were obtained from The Roslin Institute, The University of Edinburgh (McGrew et al. 2004; Davey et al. 2006). Eggs were incubated at 37 °C and staged according to the embryonic day of development (E), and by using the developmental tables of Hamburger and Hamilton (Hamburger and Hamilton 1951). For each combination of grafting experiment (chick^GFP^-*talpid^3^*; *talpid^3^*-chick^GFP^; chick^GFP^-wild type), the neural tube adjacent to somites 2–6 inclusive (and its associated neural crest) was microsurgically removed from the host embryos at embryonic day E1.5 and replaced with equivalent stage-matched tissue, as previously described (Burns and Le Douarin 2001; Delalande et al. 2015). Following grafting, eggs were returned to the incubator, and embryos allowed to develop to the appropriate stage. To ascertain the genotype of the Talpid^3^ neural tube transplant onto a GFP host embryo, the remaining tissues of the donor were used for DNA extraction, PCR amplification and sequencing of the region flanking the a366 mutation on exon7 using the following primers: Forward: CATTAGCTCTGCCGTCAACA Reverse: GGTAGGCAGACCACTGGAAG (Fig. 5) (Davey et al. 2006). A total of 30 GFP neural tube grafts into TALPID3 hosts were made, of which 2 homozygote *talpid^3^* mutant hosts were fixed for analysis at E6.5. A total of 5 *talpid^3^* neural tube grafts onto GFP hosts were made, of which 1 chimera with a homozygote *talpid^3^* mutant donor neural tube was identified after genotyping and fixed for analysis at E7.5.

### Mouse conditional knockout

Animals used for this study were maintained and the experiments were performed in accordance with local approvals and the UK Animals (Scientific Procedures) Act 1986 under licence from the Home Office (PPL70/7500). *Talpid3* ^*f/f*^ mice were acquired from Prof. Malcolm Logan (King’s College London). *Wnt1-cre;R26R-YFP/YFP* mice (Srinivas et al. 2001; Druckenbrod and Epstein 2005), in which NCC express yellow fluorescent protein (YFP), were crossed to *Talpid3* ^*f/f*^ to generate a neural crest conditional knockout of *talpid^3^*. Subsequent *Wnt1:Cre; Talpid3* ^*flox/flox*^ mouse tissues were examined at postnatal day 0 (P0).

### Human embryonic and fetal material

Human material was sourced *via* the Joint MRC/Wellcome Trust Human Developmental Biology Resource (HDBR) under informed ethical consent with Research Tissue Bank ethical approval (08/H0712/34+5 and 08/H0906/21+5) (Gerrelli et al. 2015). Staging of embryos was carried out according to the Carnegie system. GI tissues from a fetus with Short-rib polydactyly and bearing a homozygous null mutation in *KIAA0586* were obtained from case II:5 family 4, as previously described in (Alby et al. 2015). Informed consent was obtained for all participating families, and the study was approved by the ethical committee of Paris Ile de France II.

### Tissue sectioning

Transverse sections were cut from whole embryos and dissected gut segments. Sections were obtained at a thickness of 6 μm from wax blocks and 10–15 μm from frozen blocks. All sections were placed on Superfrost Plus microscope slides (BDH Laboratories).

### Immunofluorescence and in situ hybridization

For labeling of chicken and human dissected GI samples, the tissues were processed for immunofluorescence as previously described (Wallace and Burns 2005; Delalande et al. 2014) or *in situ* hybridization (Burns and Delalande 2005). For immunofluorescence, samples were incubated with primary antibodies listed in Table 1, followed by washes and incubation with fluorescently tagged secondary antibodies listed in Table 2. Samples were stained for 10 min with DAPI, before being mounted under a coverslip using Vectashield mounting medium (Vector Laboratories).

**Table 1.** Primary antibodies for immunohistochemistry studies.

| PRIMARY ANTIBODY | CONCENTRATION | COMPANY |
| --- | --- | --- |
| Mouse anti-TuJ1 | 1:500 | Covance (MMS-435P) |
| Mouse anti-CS56 | 1:2000 | SIGMA (C8350) |
| Mouse anti-Collagen IX | 1:2 | DSHB (2B9) |
| Mouse anti-SMA | 1:400 | Dako (M0851) |
| Chicken anti-GFP | 1:500 | Abcam (ab13970) |
| Chicken anti-Ncad | 1:5 | DSHB (6B3) |
| Rabbit anti-SHH | 1:400 | Santa Cruz (sc-6149) |
| Rabbit anti-PTCH | 1:100 | EMD Millipore (06-1102) |
| Rabbit anti-IFT88 | 1:400 | Thermofisher (PA56997) |
| Pan Cytokeratin-488 | 1:20 | eBioscience (53-9003-82) |
| Rabbit anti-Arl13b | 1:50 | ProteinTech (17711-1-AP) |

**Table 2.** Secondary antibodies for immunohistochemistry studies.

| SECONDARY<br>ANTIBODY | ALEXA<br>FLUOR | CONCENTRATION | COMPANY |
| --- | --- | --- | --- |
| **Goat anti-rabbit** | 488 | 1:500 | Invitrogen |
| **Goat anti-mouse** | 488 | 1:500 | Invitrogen |
| **Anti-chicken** | 488 | 1:500 | Abcam |
| **Anti-mouse** | 568 | 1:500 | Invitrogen |
| **Anti-rabbit** | 568 | 1:500 | Invitrogen |
| **Anti-mouse** | 647 | 1:500 | Invitrogen |
| **Anti-rabbit** | 647 | 1:500 | Invitrogen |

### Confocal microscopy, cell counting and statistical analysis

Tissues were imaged using confocal microscopy (Zeiss LSM 710 confocal microscope). Images of gut sections double immunostained with relevant tissue antibodies and DAPI. Equivalent fields of relevant sections were examined in controls, mutants and/or chimeras (minimum n=3). Cells were counted and tissue thickness measured using the Image-J Fiji cell counter plugin and measuring tool respectively. Data was plotted to a histogram or interval plot. The results for all samples were normalized to baseline. The outcome was set as the percentage of the difference between the baseline and the experimental conditions. Multiple linear regression was used to study the effects of tissue, species and gene mutation simultaneously. This allowed us to assess the relative contribution of each predictor to the total variance that our model explained. Interaction terms were also considered with stricter p value cut offs. The models were validated using the adjusted R squared and by performing residual analysis. To assess possible tissue rescue in the GFP>*taplid*^*3*^ transplantation experiment, the non-parametric Spearman’s Rho test was used to measure the correlation between control, chimera and mutant.

### Neural tube culture and scanning electron microscopy

Migrating vagal NCC for scanning electron microscopy imaging were obtained using *in vitro* neural tube cultures as previously described (Delalande et al. 2008). Samples were then fixed overnight in 2% glutaraldehyde, 2% paraformaldehyde in 0.1 M phosphate buffer, pH7.4, at 4°C, post-fixed in 1% OsO4/1.5% K4Fe(CN)6 in 0.1 M phosphate buffer at 3°C for 1.5 hr. After rinsing with 0.1 M phosphate buffer and distilled water, specimens were progressively dehydrated to 100% ethanol, then washed once in acetone. The samples were then critical point dried using CO2 and mounted on aluminium stubs using sticky carbon taps. Samples were then coated with a thin layer of Au/Pd (2 nm thick) using a Gatan ion beam coater and imaged with a Jeol JSM-6480LV high-performance, Variable Pressure Analytical Scanning Electron Microscope.

### Mouse skeletal staining

*Wnt1:Cre; Talpid3* ^*fl/fl*^ P0 mice and control littermates were fixed in 90% ethanol, then skinned and eviscerated. Staining with Alcian Blue (0.05%) was performed in 70% ethanol with 20% acetic acid, followed by staining with Alizarin Red (0.15%) in 1% KOH. Soft tissue was cleared with 1% KOH with 20% glycerol and skeletons were stored in 80% glycerol.

## Acknowledgements

The authors would like to thank Dr Dagan Jenkins for the IFT88 antibody, Mark Turmaine for assistance with SEM, Dr Kevin Lee and Dr Erwin Pauws for assistance with cartilage and bone staining as well as help with the phenotype analysis. We also thank Dr Jan Soetaert and Dr Belén Martín-Martín for help with microscopy. M.D. and the TALPID3 flock are supported by an Institutional Strategic Grant (ISP) to The Roslin Institute from the BBSRC. N.N. is supported by a Bolyai Fellowship and Hungarian Science Foundation NKFI grant (124740). CM is supported by Guts UK (Derek Butler Fellowship). We are grateful to the French Society of Fetal Pathology (SoFFoet) for participating in the study.

We acknowledge the NIHR Great Ormond Street Hospital Biomedical Research Centre which supports all research at Great Ormond Street Hospital NHS Foundation Trust and UCL Great Ormond Street Institute of Child Health. The views expressed are those of the authors and not necessarily those of the NHS, the NIHR or the Department of Health.

---

**Supplementary Figure S1:**
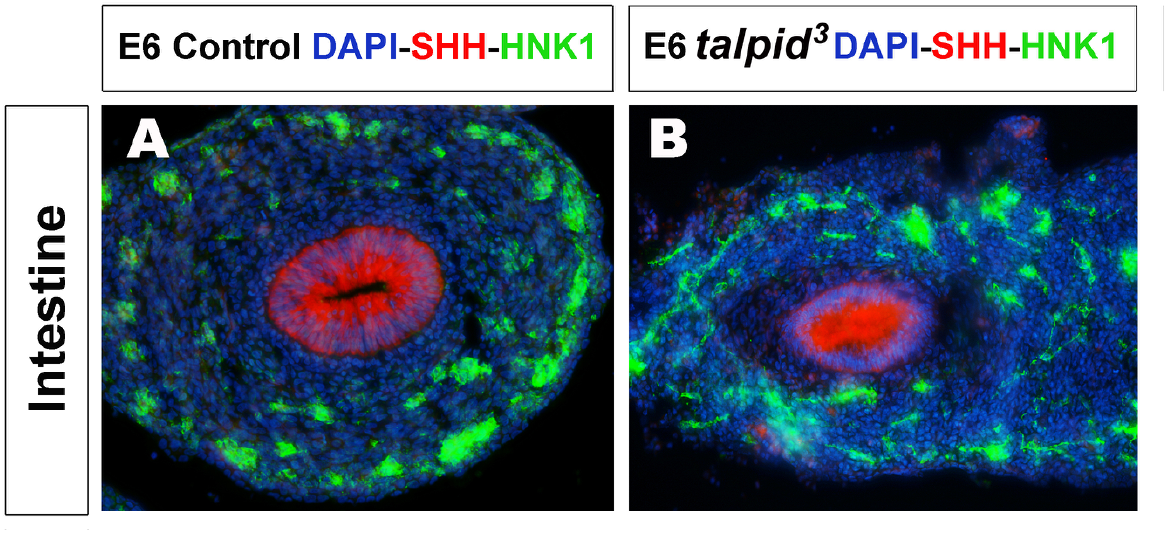
The protein Sonic Hedgehog is expressed in both control and *talpid^3^* intestine. (A,B) Immunofluorescent staining of intestine with SHH and HNK1 in (A) E6.5 control and (B) *talpid^3^* mutant embryo. SHH (red) is expressed in the epithelium of both samples. Scattered HNK1+ ENCC (green) are observed in the intestine of *talpid^3^* mutant embryo.

**Supplementary Figure S2:**
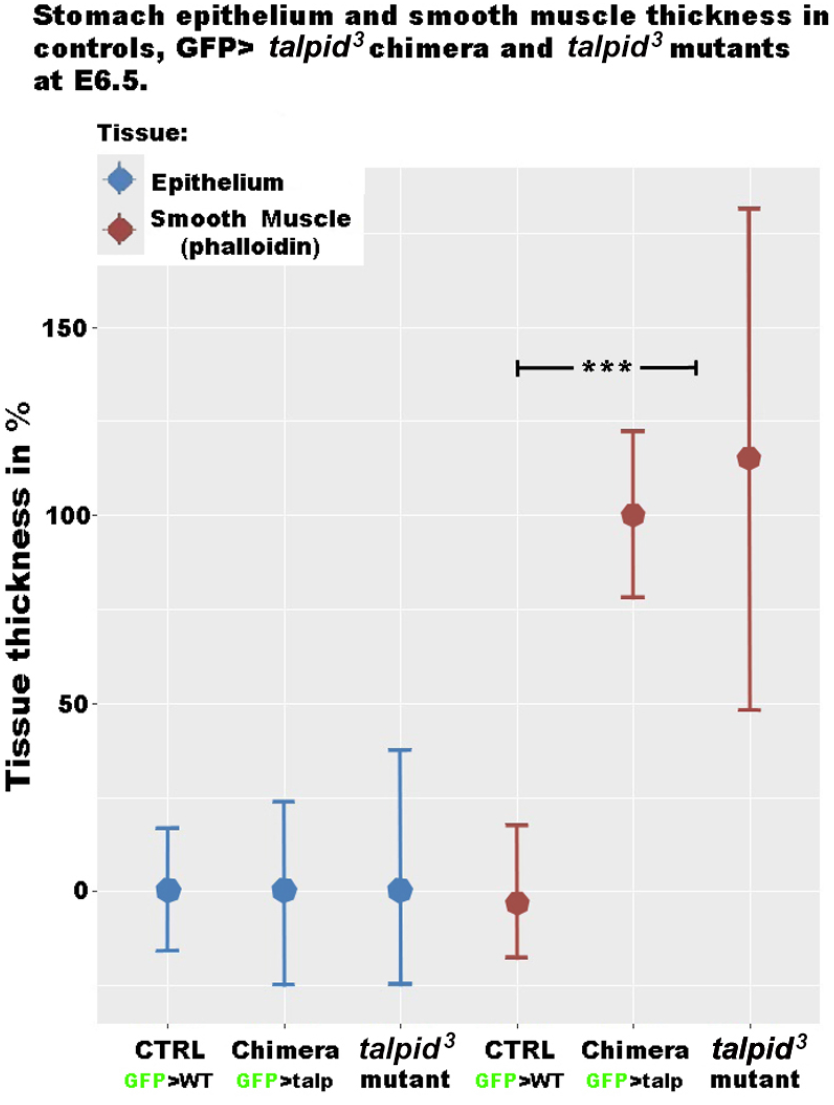
Transplantation of wild type ENCCs does not rescue the muscle phenotype in GFP>*talpid^3^* chimeric E6.5 embryo.. Measurements of epithelium (DAPI) and smooth muscle (Phalloidin) thickness in stomach and intestine sections were normalised to baseline. At E6.5 there was no statistical difference between the thickness of the epithelium of controls versus GFP>*talpid^3^* chimera or *talpid^3^* mutants. Thickness of the smooth muscle, as measured by phalloidin was increased 100% in the chimera and 115% in the mutant compared to GFP>wild type control transplant. Measurements of phalloidin thickness in GFP>*talpid^3^* chimera and mutant were statistically equivalent and they were both statistically significant from the baseline control (*** *p<0*.*001*).

**Supplementary Figure S3:**
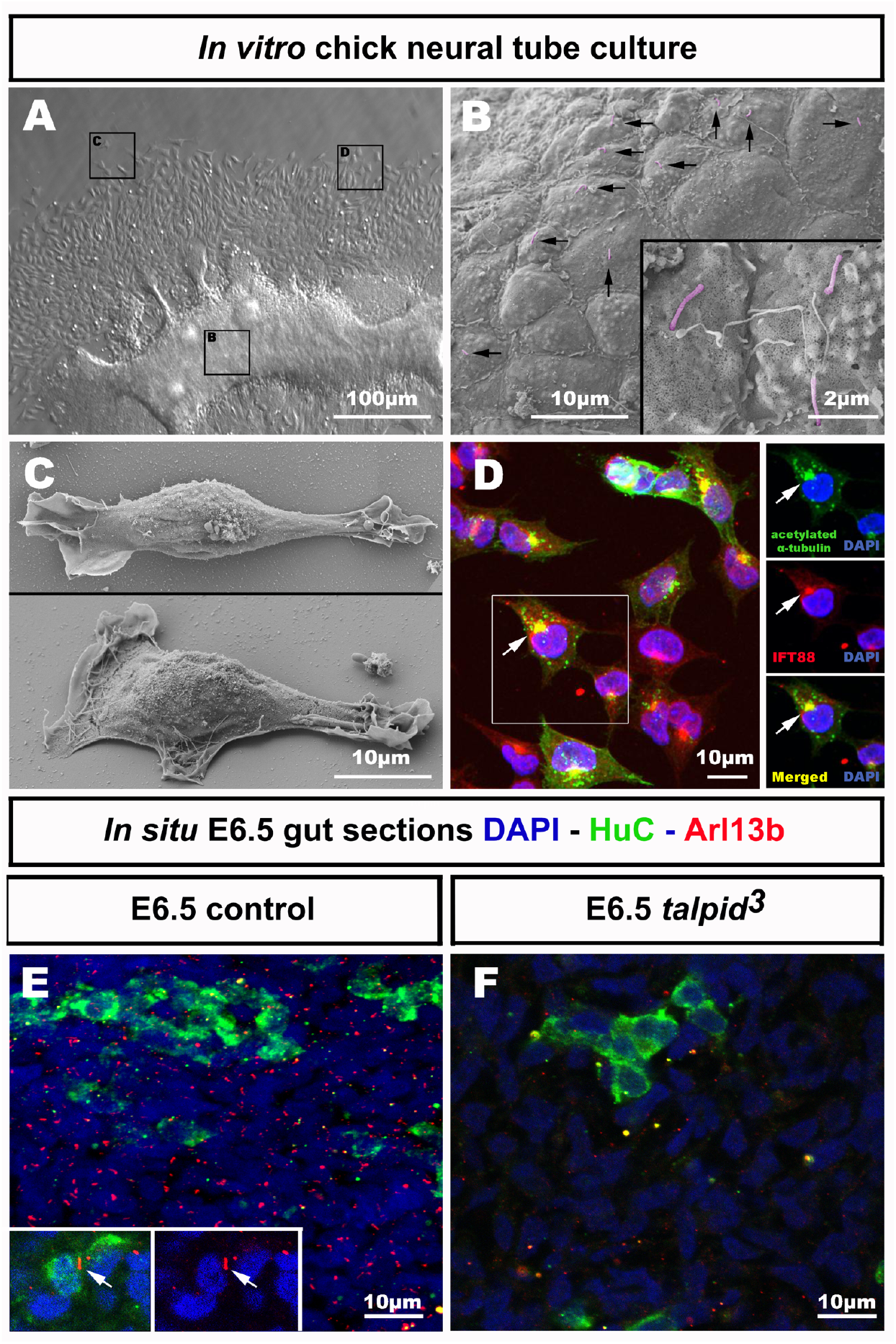
Migrating ENCCs do not extend a primary cilium. (A-C) Scanning electron microscopy of neural tube culture and migrating vagal neural crest after 18h culture. (A) Low magnification of picture shows neural tube and migrating vagal neural crest spreading out. (B) Close up of the neural tube shows primary cilia on cells (pseudocolored in purple, black arrows). (C) Close up of migrating vagal NCC shows no primary cilium. (D) Immunofluorescent staining for acetylated α-tubulin (green) and intraflagellar transport protein 88 (red) shows colocalisation at the centrosome and no primary cilium on migrating vagal NCC (white arrow). (E) *In vivo* staining for the primary cilia with Arl13b on E6.5 gut sections shows signal in all mesenchymal gut cells. In migrating HuC^+^ (green) ENCCs,11% show an extended primary cilium (inset; 10 out of 85 HuC^+^ cells counted, n=7 sections). (F) No primary cilia staining is observed in any cell type on *talpid^3^* E6.5 gut section.

**Supplementary Figure S4:**
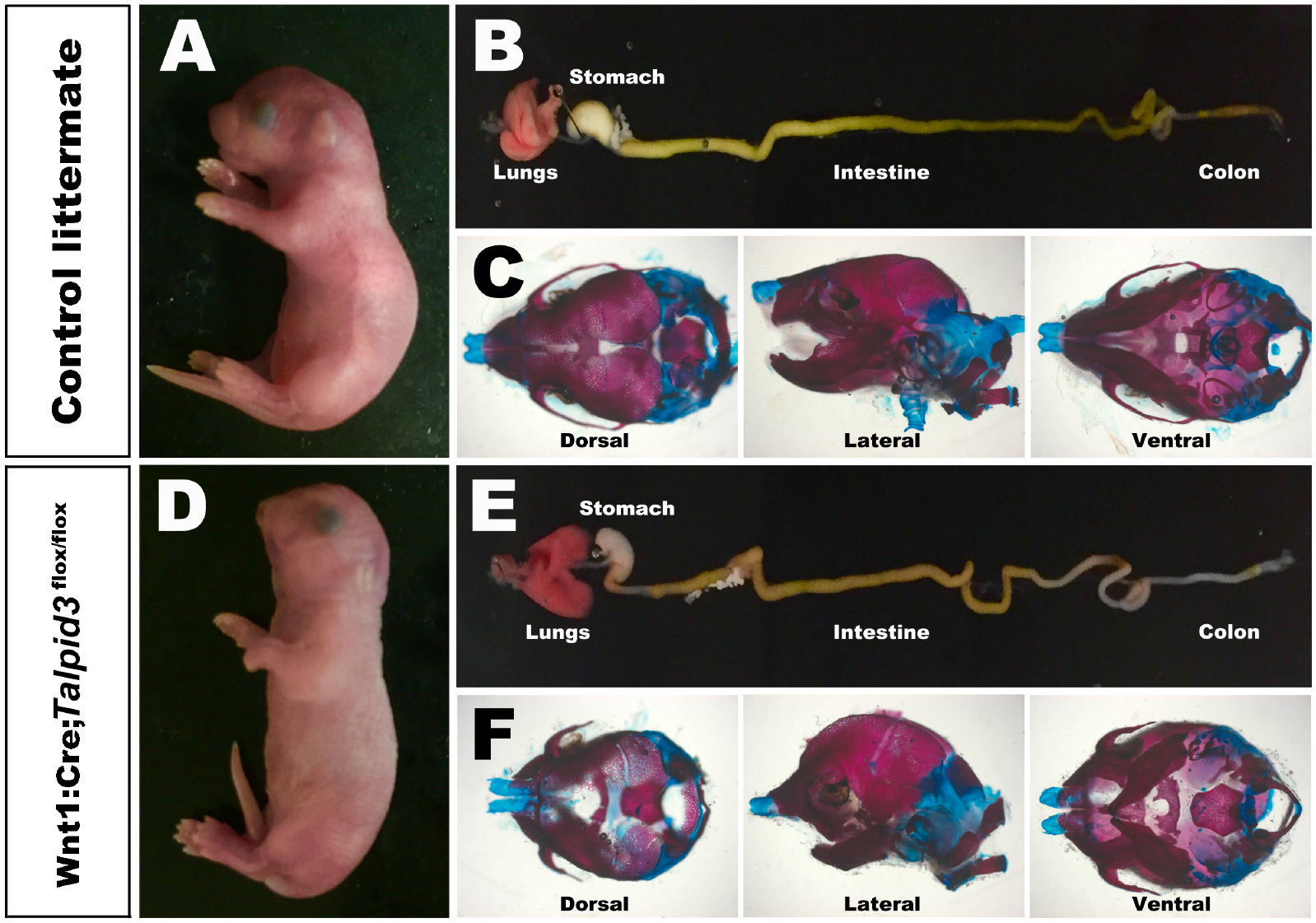
Cranial and GI tract phenotypes of *Wnt1:Cre;Talpid3*^*fl/fl*^ P0 mice and control littermate. (A,D) Gross phenotype of (A) control littermate and (D) *Wnt1:Cre;Talpid3*^*fl/fl*^ P0 pups. (B,E) Gross morphology of the dissected gastrointestinal tract and lungs of (B) control littermate and (E) *Wnt1:Cre;Talpid3*^*fl/fl*^ P0 pups. Mutant littermates show grossly normal GI tract, red coloured uninflated lungs as well as craniofacial abnormalities such as cleft pallet, short snot and brachycephaly. (C,F) Skeletal staining of the skulls of (C) control littermate and (F) *Wnt1:Cre;Talpid3*^*fl/fl*^ P0 pups. Red indicates bone and blue indicates cartilage. (F) *Wnt1:Cre;Talpid3*^*fl/fl*^ skull shows frontonasal hypoplasia, hypoplastic NCC derivatives, micrognathia, facial cleft-partitioning of the nasal cartilage and underdeveloped sagittal suture.

**Supplementary Figure S5:**
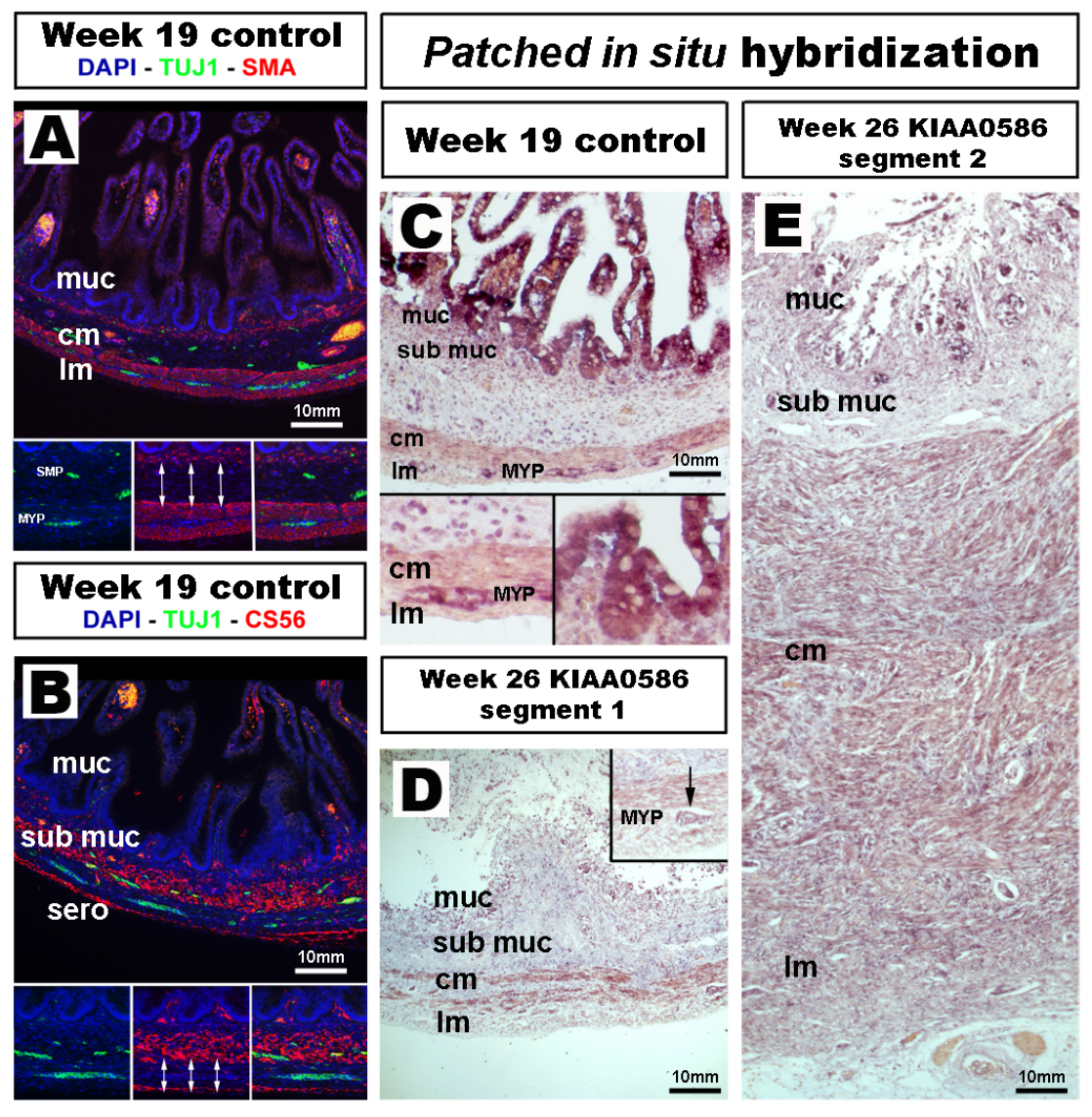
Expression of SMA, Tuj-1, CS56 and *patched* in week 19 control human gut and *patched* in situ hybridization in 26 week KIAA0586 gut. (A,B) Control intestine shows normal neuromuscular patterning and CS56 expression at week 19. (C-E) *In situ* hybridization against *patched* in week 19 control and *KIAA0586* mutated tissue. (C + insets) Week 19 control intestine shows strong expression of *patched* in the epithelium, in the myenteric plexus of the ENS and to a lesser extend in the smooth muscle. (D) “Segment 1” shows *patched* expression in the myenteric plexus (inset) and the muscle layers. (E) “Segment 2” shows a diffuse *patched* expression throughout the section. muc: mucosa; sub muc: submucosa; cm: circular muscle; lm: longitudinal muscle; MYP: myenteric plexus.

**Supplementary video S1:**
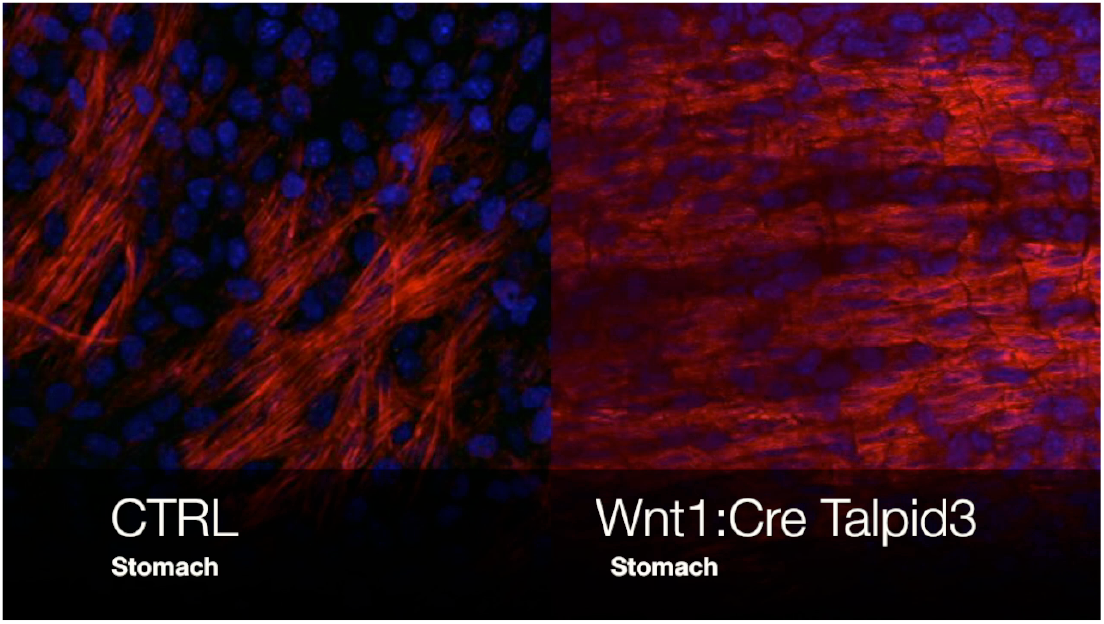
Animated confocal Z stack of whole mount immunofluorescent staining for SMA smooth muscle actin (red) and TuJ-1 Neuron-specific class III beta-tubulin (green) and DAPI (Blue) on stomach of control P0 mice littermate (left) and *Wnt1:Cre;Talpid3*^*fl/fl*^ (right).

**Supplementary video S2:**
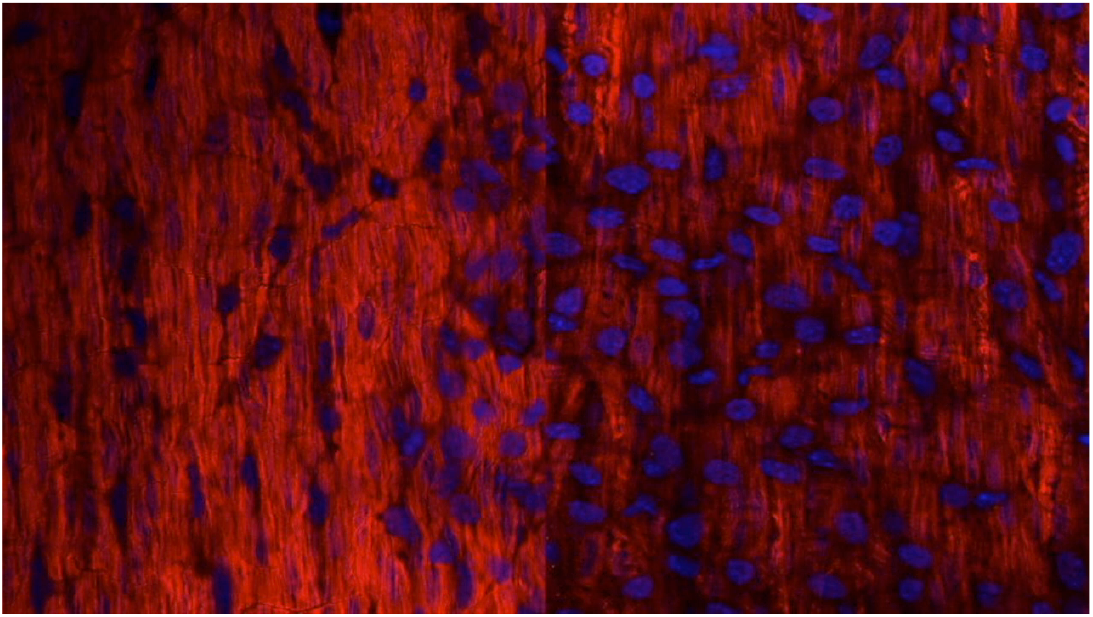
Animated confocal Z stack of whole mount immunofluorescent staining for SMA smooth muscle actin (red) and TuJ-1 Neuron-specific class III beta-tubulin (green) and DAPI (Blue) on intestine of control P0 mice littermate (left) and *Wnt1:Cre;Talpid3* ^*fl/fl*^ (right).

## Notes

### Competing Interest Statement

The authors have declared no competing interest.

